# Unification of optimal targeting methods in Transcranial Electrical Stimulation

**DOI:** 10.1101/557090

**Authors:** Mariano Fernandez-Corazza, Sergei Turovets, Carlos Muravchik

## Abstract

One of the major questions in high-density transcranial electrical stimulation (TES) is: given a region of interest (ROI), and given electric current limits for safety, how much current should be delivered by each electrode for optimal targeting? Several solutions, apparently unrelated, have been independently proposed depending on how “optimality” is defined and on how this optimization problem is stated mathematically. Among them, there are closed-formula solutions such as ones provided by the least squares (LS) or weighted LS (WLS) methods, that attempt to fit a desired stimulation pattern at ROI and non-ROI, or reciprocity-based solutions, that maximize the directional dose at ROI under safety constraints. A more complete optimization problem can be stated as follows: maximize directional dose at ROI, limit dose at non-ROI, and constrain total injected current and current per electrode (safety constraints). To consider all these constraints (or some of them) altogether, numerical convex or linear optimization solvers are required. We theoretically demonstrate in this work that LS, WLS and reciprocity-based closed-form solutions are particular solutions to the complete optimization problem stated above, and we validate these findings with simulations on an atlas head model. Moreover, the LS and reciprocity solutions are the two opposite cases emerging under variation of one parameter of the optimization problem, the dose limit at non-ROI. LS solutions belong to one extreme case, when the non-ROI dose limit is strictly imposed, and reciprocity-based solutions belong to the opposite side, i.e., when this limit is loose. As we couple together most optimization approaches published so far, these findings will allow a better understanding of the nature of the TES optimization problem and help in the development of advanced and more effective targeting strategies.

## 1 Introduction

Transcranial electrical stimulation (TES) is an emerging therapy for the treatment of neuropsychiatric conditions such as depression (Kalu et al., 2012), Parkinson’s disease (Boggio et al., 2006), anxiety and chronic pain (Mori et al., 2010). Research has also demonstrated that TES can be a valuable therapeutic tool in epilepsy (Yook et al., 2011), stroke rehabilitation (Schlaug et al., 2008), and other neurological and psychiatric conditions (Brunoni et al., 2013). It has also been extensively studied in the context of enhancing cognitive skills such as memory and learning (Berryhill and Jones, 2012; Nitsche et al., 2003). This technique may become eventually an alternative for psychoactive drugs, as it can be more selective than drugs by targeting specific regions of interest in the brain with minimal adverse side effects and it does not affect the entire brain indiscriminately. Even without producing direct neuronal firing, TES application is capable to modify cortical excitability (Nitsche and Paulus, 2000; Priori et al., 1998) as well as brain rhythms and networks (Lang et al., 2005; Priori, 2003), and this is why the method is also termed Transcranial Electrical Neuromodulation (TEN). Despite recent advances, there are ongoing debates on the clinical effectiveness of TES (Antal et al., 2015; Horvath et al., 2015, 2014) addressing many issues to be still resolved, in particular, substantial inter-subject response variability (Batsikadze et al., 2013; Wiethoff et al., 2014). The general idea is that optimal targeting protocols and the use of subject-specific accurate head models might enhance rigor and reproducibility in TES (Bikson et al., 2018). Because the goal is to stimulate the brain, TES is also termed Transcranial Brain Stimulation (TBS). If direct or alternating currents are used, TES is termed transcranial direct current stimulation (tDCS) or transcranial alternating current stimulation (tACS) respectively.

In TES, electric currents are applied to two or more electrodes placed on the scalp. If the number of electrodes is larger than 2, it is called multi-electrode TES. If it is even larger, being for instance 32, 64, 128 or 256 like typically arranged in high channel count electroencephalography (EEG), it is known as high-density TES. A list of electric current levels applied to the head at each electrode is known as a current injection pattern. A current injection pattern generates an electric field (or current density) map on the brain, which can be considered as the actual dose power in TES. Given low frequencies of the injected currents, the quasi-static approximation of the Maxwell equations governs the physics involved in the computation of this map, which is known as the TES forward problem (FP). TES FP is typically solved numerically using finite element method (FEM) (Datta et al., 2013), boundary element method (BEM) (Goncalves et al., 2003) or finite difference method (FDM) (Turovets et al., 2014).

The inverse problem (IP) goal in high-density TES is to determine how much current should be delivered by each electrode for optimal targeting a particular region of interest (ROI) within the brain, i.e., determining optimal (in some sense) current injection patterns for a given ROI. When solving the TES inverse problem, one should address a trade-off between maximizing the electric field at the ROI and minimizing it at the non-ROI and, at the same time, limit values of applied currents for safety. The two common limits are: total injected current, which can be thought as a fixed budget, and maximum current per electrode. Depending on the optimality criteria, several schemes have been proposed leading to different optimal solutions.

**Least Squares (LS) and Weighted-LS (WLS)** are the simplest and most typical optimization methods. The LS solution derives from minimizing a second-order error between the resulting and the desired electric field (or current density) profiles at a particular domain of interest Ω (typically, the ROI is included in Ω, and ROI much smaller than Ω). Usually, Ω is the gray matter or the entire brain, where a desired electric field is set to some desired profile (including directions and intensities) at the ROI and zero at the non-ROI of Ω (Dmochowski et al., 2011; Guler et al., 2016), but also Ω can be the entire head (Fernández-Corazza et al., 2016). WLS is equivalent to LS with the addition of a weight matrix that, for instance, can control intensity-focality trade-off (Dmochowski et al., 2011) or incorporate additional *a-priori* knowledge (Ruffini et al., 2014). If no current injection limits are imposed or they are too loose to be neglected, the LS or WLS solutions can be presented by a well-known closed formula (Dmochowski et al., 2011; Fernández-Corazza et al., 2016; Salman et al., 2016). One option to account for the total current budget constraint without the need of numerical solvers is to apply a scaling factor to the closed formula (as in (Dmochowski et al., 2017; Fernández-Corazza et al., 2016)) and, as we show later, the solution is still optimal. Another option is to consider the total and per electrode current limits and solve the problem using a numerical optimization algorithm such as LASSO (Dmochowski et al., 2011), MATLAB convex optimization (Dmochowski et al., 2011) or genetic algorithms (Otal et al., 2016; Ruffini et al., 2014). The LS based optimization was also earlier formulated in the context of multichannel TMS (Ilmoniemi et al., 1999).

**Constrained directional maximization** of the electric field (or current density) intensity at the target along a predefined and desired orientation is another optimization approach. In this approach, the functional to maximize is linear, thus it requires some limiting restrictions or constraints (such as total current injection budget) to get finite solutions. It can be numerically solved with convex optimization packages such as CVX (Grant and Boyd, 2014). Dmochowski et al 2011 (Dmochowski et al., 2011) solved a reduced version of this problem for the electrical field considering only the total current limit (Eq. 17 in (Dmochowski et al., 2011)), whereas Guler et al 2016 (Guler et al., 2016) and Wagner et al 2016 (Sven Wagner et al., 2016) considered a more complete optimization problem including additional constraints of an upper bound for the electric field at the non-ROI cortex and a per-electrode current limit.

**Reciprocity-based** optimization solutions are based on the reciprocity theorem (Malmivuo and Plonsey, 1995; Rush and Driscoll, 1969). According to this theorem, the optimal (in terms of maximizing directional intensity) solution to target a given ROI along a given direction, can be related to the EEG forward projection to the scalp of source dipoles artificially placed at the same ROI in the direction of interest (Cancelli et al., 2016; Dutta and Dutta, 2013; Fernández-Corazza et al., 2016; Rush and Driscoll, 1969; Salman et al., 2016). The “EEG forward projection” means the electric potential on the scalp produced by the neuronal sources (typically modelled as electrical dipoles), which is known as the EEG forward problem. As the reciprocity-based solutions are optimal and not iterative, they can be also considered “closed-formula” solutions. One reciprocity approach is to concentrate the electric current sources and sinks as close as possible to the “poles” of the EEG forward projection (Fernández-Corazza et al., 2016, 2015; Guhathakurta and Dutta, 2016). In our previous work, we mathematically demonstrated that this strategy maximizes the directional electric field at the ROI given a fixed current injection budget (Fernández-Corazza et al., 2016). Another approach is setting the current injection pattern proportionally to the EEG forward projection, either directly or after applying a Laplacian filter (Cancelli et al., 2016; Dutta and Dutta, 2013), though we found that its performance was not better in any of the tested metrics compared to other approaches (Fernandez-Corazza et al., 2017).

In this work, we link these three apparently unrelated optimization approaches and some of their variants resulting in a unified approach that couples together almost all optimization approaches described so far (see Section 5.7 for a list of included and not included approaches in this unified approach). As far as we know, the links we present here have not been fully noticed previously. First, we analytically demonstrate that the **constrained maximization** of directional intensity approach can be expressed by a **WLS** closed formula when considering a strict upper bound for the non-ROI intensity. Consequently, we show that the unrestricted closed formula **LS** or **WLS** solution, scaled in the way that the total budget is exploited, is optimal in some sense and it doesn’t require numerical solvers. Second, we analytically demonstrate that the **constrained maximization** solution equals to a **reciprocity-based** solution (one source and one sink at the EEG poles) when the non-ROI intensity bound is too loose. This idea was already demonstrated in our previous work, but we now theoretically link it to the numerical solution of the more general optimization problem. Moreover, by imposing different current limit per electrode bounds, the **constrained maximization** numerical solutions resemble some variants of the **reciprocity** solutions of our previous work (Fernández-Corazza et al., 2016) that use more than one electrode as sources or sinks. This implies that the **reciprocity-based** solutions concentrating the sources and sinks in clusters as close as possible to the EEG poles are by no means ad-hoc, but optimal in sense of maximizing directional intensity of stimulation at ROI. Lastly, if the non-ROI electric field bound is neither too loose nor too strict, the full **constrained maximization** problem can be solved numerically using a convex optimization package such as CVX as in (Guler et al., 2016). In this middle range, the solutions show a smooth transition from closed-form **WLS** to closed-form **reciprocity** as the non-ROI electric field upper bound gets looser (larger). Within this middle range, we also empirically found that the **LS** with total current limit constraint numerical solution (solved using LASSO by Dmochowski et al in (Dmochowski et al., 2011)) is also included within the framework of the **constrained maximization** approach.

We show simulation results when the brain is considered as the domain of interest Ω (typical), and also when using the full head domain as Ω. The latter, is a novel approach first used in our previous work (Fernández-Corazza et al., 2016): as current budget is limited, it makes sense to minimize the electric field at the rest of the whole head and not just the rest of the brain (or cortex) just not to “waste” the budget.

Other less common optimization approaches have been also proposed in the literature that are not considered in this work. One of them is beamforming or Linearly Constrained Minimum Variance (LCMV) (Dmochowski et al., 2011; Fernández-Corazza et al., 2016). This approach implies that electric field at the ROI or target is totally collinear with desired targeting orientation. Similarly to LS or WLS, it has a closed formula solution when no current limits are considered. Another approach is maximizing module of the electric field at ROI instead of directional intensity (Sadleir et al., 2012). This problem, although it has great interest for the application, is much more difficult to solve as it is not convex nor linear. The authors attempted to solve it using the interior point optimization algorithm, but they concluded that there is no guarantee that the solution they found is a global optimum or even unique due to the complex nature of this optimization problem (Sadleir et al., 2012).

## 2 TES forward problem

Due to the low frequencies involved, the TES forward problem (FP) is governed by quasi-static Maxwell equations, the Poisson equation for the electric potential 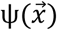 in the head volume with Neumann boundary conditions (Frank, 1952; Jackson, 1975). It is typically solved using the Finite Element Method (FEM) (Kwon and Bang, 2000; Silvester and Ferrari, 1994), where the whole head is meshed into *N*_*H*_ elements, usually tetrahedrons, and *P* nodes. The details of the FEM FP formulation in TES can be found elsewhere (Laakso et al., 2016; Ruffini et al., 2014; Vauhkonen et al., 1999; Windhoff et al., 2013). Note that the FEM FP is equivalent to the Electrical Impedance Tomography (EIT) FP, and thus, EIT literature also details the same FEM formulation (Abascal et al., 2008; Fernández-Corazza et al., 2013; Lionheart et al., 2004; Wang et al., 2009).

The boundary conditions differ in approximation of pointwise or distributed electrodes. In the last case, they are modelled using the complete electrode model (CEM) (Hyvönen, 2004). In any case, FEM results in a linear problem **Kv = f**, where **K** is called *stiffness matrix* and accounts for geometry, conductivity or conductivity map of each tissue, and electrode contact impedances (if using CEM); **v** is the unknown electric potential at each mesh node of the head and electrodes (if using CEM (Lionheart et al., 2004)), and **f** is an independent vector accounting for the electric sources and sinks (in TES, the applied currents or, equivalently, the current injection pattern). Once the system of linear equations above is solved for **v**, for instance using preconditioned conjugate gradients (Barrett et al., 1994) or biconjugate stabilized gradient (van der Vorst, 1992), the electric field 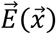 can be easily computed at each element by: 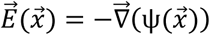, where 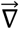 is the gradient operator.

## 3 Unification of optimization approaches

Table I summarizes all different optimization methods that are covered by the unified approach. The solutions to all these problems of Table I can be found as particular solutions of the complete constrained maximization approach of Guler et al 2016 (Guler et al., 2016), which we describe next in section 3.1. In sections 3.2 and 3.3 we link the solution of this approach to the other approaches, specifically to LS/WLS and reciprocity-based solutions.

**Table I:** Summary of all covered approaches in the unified framework. Thus, all approaches listed here can be though as particular versions of the complete approach stated in eq. (3).

| Approach | Formulation<sup>1</sup> | Solution | References |
| --- | --- | --- | --- |
| **LS** | $\hat{\mathbf{i}} = \underset{\mathbf{i}}{\operatorname{argmin}}\{(\mathbf{e} - \mathbf{Ti})^T(\mathbf{e} - \mathbf{Ti})\}$ | $\hat{\mathbf{i}} = (\mathbf{T}^T\mathbf{T})^{-1}\mathbf{T}^T\mathbf{e}$ | - Dmochowski et al 2011, eq (6) (Dmochowski et al., 2011)<br>- Salman et al 2015, eq (8) (Salman et al., 2016) |
| **Scaled LS** | $\hat{\mathbf{i}} = \underset{\mathbf{i}}{\operatorname{argmin}}\{(\mathbf{e} - \mathbf{Ti})^T(\mathbf{e} - \mathbf{Ti})\}$ | $\hat{\mathbf{i}} = k(\mathbf{T}^T\mathbf{T})^{-1}\mathbf{T}^T\mathbf{e}$ ,<br>where $k$ is a scaling constant such that $\ \hat{\mathbf{i}}\ _1 = 2i_{max}$ | - Fernandez-Corazza et al 2016, eq (4) and “Least Squares” subsection (Fernández-Corazza et al., 2016)<br>- Dmochowski et al 2017, eq (4) (Dmochowski et al., 2017) |
| **WLS** | $\hat{\mathbf{i}} = \underset{\mathbf{i}}{\operatorname{argmin}}\{(\mathbf{e} - \mathbf{Ti})^T\mathbf{\Gamma}(\mathbf{e} - \mathbf{Ti})\}$ | $\hat{\mathbf{i}} = (\mathbf{T}^T\mathbf{\Gamma}\mathbf{T})^{-1}\mathbf{T}^T\mathbf{\Gamma}\mathbf{e}$ | |
| **Total current constrained LS/WLS** | $\hat{\mathbf{i}} = \underset{\mathbf{i}}{\operatorname{argmin}}\{(\mathbf{e} - \mathbf{Ti})^T\mathbf{\Gamma}(\mathbf{e} - \mathbf{Ti})\}$<br>$s. t. \ \hat{\mathbf{i}}\ _1 < 2i_{max}$ | Numerical, LASSO algorithm | - Dmochowski et al 2011, eq (10) (WLS) (Dmochowski et al., 2011)<br>- Dmochowski et al 2017, eq (8) (LS) (Dmochowski et al., 2017) |
| **Fully constrained WLS** | $\hat{\mathbf{i}} = \underset{\mathbf{i}}{\operatorname{argmin}}\{(\mathbf{e} - \mathbf{Ti})^T\mathbf{\Gamma}(\mathbf{e} - \mathbf{Ti})\}$<br>$s. t. \ \hat{\mathbf{i}}\ _1 < 2i_{max}$<br>and $\hat{\mathbf{i}}_{min} \leq \hat{\mathbf{i}} \leq \hat{\mathbf{i}}_{max}$ | Numerical, MATLAB convex iterative algorithm<br>Numerical, genetic algorithm | - Dmochowski et al 2011, eq (11) (Dmochowski et al., 2011)<br>- Ruffini et al 2014, eq (1) (Ruffini et al., 2014)<br>*Note: They use different weighting matrices $\mathbf{\Gamma}$* |
| **Simple maximizing intensity** | $\hat{\mathbf{i}} = \underset{\mathbf{i}}{\operatorname{argmax}}(\mathbf{e}^T\mathbf{Ti})$ ,<br>$s. t. \ \hat{\mathbf{i}}\ _1 < 2i_{max}$ | Numerical, linear programming | - Dmochowski et al 2011, eq (17) (Dmochowski et al., 2011)<br>- Cancelli et al 2016, “Model based intensity, 2 electrodes” (Cancelli et al., 2016) |
| **Complete maximizing intensity** | $\hat{\mathbf{i}} = \underset{\mathbf{i}}{\operatorname{argmax}}(\mathbf{e}^T\mathbf{\Gamma}\mathbf{Ti})$ ,<br>$s. t. \mathbf{i}^T\mathbf{T}^T\mathbf{\Gamma}_{non-roi}\mathbf{Ti} \leq \alpha$<br>and $\ \hat{\mathbf{i}}\ _1 < 2i_{max}$<br>and $\hat{\mathbf{i}}_{min} \leq \hat{\mathbf{i}} \leq \hat{\mathbf{i}}_{max}$ | Numerical, CVX Matlab convex optimization package | - Guler 2016, eqs (5) and (6) (Guler et al., 2016)<br>- Wagner et al 2016, problem (P) (*without current limit per electrode constraint*) (Sven Wagner et al., 2016) |
| **One-to-one reciprocity** | Simulate a dipole at the target, compute the resulting electric potential on the scalp. | Closed-form, pick as source and sink the electrodes with maximum potential difference (i.e., the EEG forward projection “poles”). | - Fernandez Corazza et al 2016, Figure 3. (Fernández-Corazza et al., 2016)<br>- Cancelli et al 2016, “Ad-hoc, voltage, sink to sink, 1x1” (Cancelli et al., 2016) |

<sup>1</sup> We use our own notation (the meaning of each symbol/letter can be found in section 3.1) that differs from the authors’ original notation.

|  |  |  |  |
| --- | --- | --- | --- |
|  |  |  | - Salman et al 2016, section 3.2 (Salman et al., 2016) |
| **Constrained reciprocity** | Simulate a dipole at the target, compute the resulting electric potential on the scalp. | Closed-form, pick as sources and sinks the electrodes with maximum potential differences (closest to the EEG forward projection “poles”) according to the maximum current limit per electrode restriction. | - Fernandez-Corazza 2016 “One source - all sinks” and “Opposite” approaches. (Fernández-Corazza et al., 2016)<br>- Cancelli et al 2016, “Ad-hoc, voltage, sink to sink, 2x2 and 4x4” (Cancelli et al., 2016) |

### 3.1 Complete constrained maximization of directional intensity approach

The complete constrained maximization approach considers the maximization of the integral of local electric field 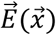 (or current density) projection onto a desired orientation 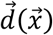 over the ROI subject to the following constraints: (i) an upper limit *α* for the electric field energy at the non-ROI in the head (or brain) domain Ω, (ii) total current limit, and (iii) current limits per electrode. The mathematical formulation can be stated as follows (Guler et al., 2016):

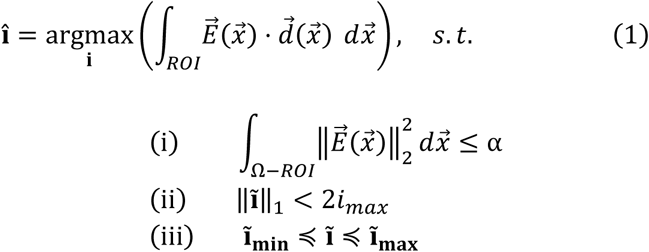

Where **i** is the unknown (*L* – 1) × 1 current injection pattern; *i*_*max*_ is the maximum total current intensity scalar; **ĩ** is the expanded current injection pattern vector of size *L* × 1 that considers all electrodes; **ĩ**_**min**_ and **ĩ**_**max**_ are the *L* × 1 minimum and maximum limits per electrode respectively; symbol ≼ means elementwise; ‖⋅‖_1_ is the *𝓁*_1_-norm (sum of absolute values of all vector components) and *L* is the number of electrodes. For *L* electrodes, there are (*L* – 1) independent current injection electrodes (pattern **i**), as the remaining electrode (the last element of expanded pattern **ĩ**) is the sum of all other currents such that total injected current is zero, i.e. (Guler et al 2016 (Guler et al., 2016), page 3, right column, last paragraph; Dmochowski et al 2011 (Dmochowski et al., 2011) restriction in Eq (10)):

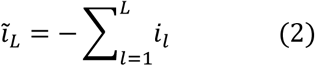

Let’s consider *L* electrodes, *N*_*H*_ total head mesh elements, and *N*_B_ brain (or cortex) mesh elements. Assume that **T**_**H**_ is the TES 3*N*_*H*_ × (*L* – 1) *transfer matrix* where each column “*l*” is the TES FP solution, i.e. electric field (or current density), at each element of the *full-head* FE mesh produced by a current injection pattern that consists of injecting the electric current at electrode *l* with last electrode *L* being the sink (or reference). Note that for *L* electrodes, there are *L* – 1 independent current injection patterns. All other patterns can be generated from this *basis* by superposition. Other bases can be used such as injecting the electric current at electrode *l* and assuming all other *L* – 1 electrodes as sinks (as used in (Fernández-Corazza et al., 2016)).

Similarly, we can define a *trimmed-to-brain* or reduced transfer matrix **T**_**B**_, where only the rows corresponding to the brain elements of the FE mesh are taken from **T**_**H**_. In what follows, either *N*_*H*_ and **T**_**H**_ or ***N***_*B*_ and **T**_**B**_ can be considered as *N* and **T** when considering full-head (subscript H) or trimmed-to-brain (subscript B) domains respectively.

The restricted maximum intensity optimization problem (1) can be stated as follows in terms of the finite element mesh (Guler et al., 2016):

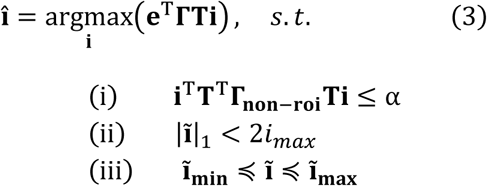

where **e** is the 3*N* × 1 desired electrical field shape, typically with non-zero values at the corresponding ROI elements (unitary oriented vectors) and zeroes at the non-ROI elements.

Volume matrix **Γ** is a diagonal 3*N* × 3*N* matrix where each element of the diagonal is the volume of the mesh element n, and **Γ**_**non-roi**_ is **Γ** but with the diagonal elements corresponding to the ROI set to zero. Matrices **Γ** and **Γ**_**non-roi**_ derive from the integration operations in Eq. (1)^2^. In the non-ROI electric field energy constraint (1.i), the integral is over the non-ROI, that’s why the elements of **Γ** corresponding to the ROI should be set to zero generating **Γ**_**non-roi**_ in (3.i). However, the ROI is typically much smaller than the non-ROI, thus **Γ**_**non-roi**_ ≈ **Γ** in sense of the norm of matrix difference. This approximation can also be interpreted as integrating constraint (1.i) in the whole domain of interest Ω (either the head or the brain), and not just at the non-ROI of Ω. The resulting difference when doing so is in order of the average intensity at the ROI times the ROI volume, over average intensity at non-ROI times non-ROI volume.

**T** in Eq. (3) is the *electric field* transfer matrix as explained before. Alternatively, one can consider the matrix product **ΣT** as the *current density* transfer matrix **T**′, where conductivity matrix **Σ** is a 3*N* × 3*N* symmetric block diagonal matrix where each 3×3 block of the diagonal is the conductivity tensor of the mesh element *n*. If piecewise isotropic media is assumed, **Σ** is a diagonal matrix, and moreover, if the trimmed-to-brain (or cortex) approach is used, and brain (or cortex) is assumed to have homogeneous and isotropic conductivity *σ*_*B*_, matrix **Σ** can be replaced by this scalar.

The earlier approach by Dmochowski et al 2011, Eq. (17) in (Dmochowski et al., 2011), is different from Eqs. (3) above in several aspects. First, non-ROI energy constraint (3.i) was not considered. Second, the pointwise sum instead of the integral was used and thus, **Γ** equals to the identity matrix in their formulation. Lastly, electric field instead of current density (as in Guler et al 2016 (Guler et al., 2016)) transfer matrix was used. We show in section 3.3 that when omitting constraint (3.i), the numerical solution of (3) is equivalent to a closed-form reciprocity-based solution, which was not noticed in both (Dmochowski et al., 2011) and (Guler et al., 2016).

### 3.2 Link between constrained directional intensity maximization and least squares approaches

In Appendix A we show that the two mathematical optimization problems below are equivalent up to a scaling constant *k*(*α*):

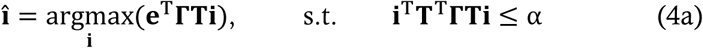

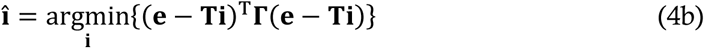

Constrained maximizing intensity problem (4a) is a bounded linear maximization problem and it is known that the solution, if not infinity or minus infinity, will lie at the boundary, i.e. at i^T^**T**^T^**ΓTi** = α (note “=” instead of “≤” sign) (Boyd and Vandenberghe, 2004). Problem (4b) is typical WLS with known analytical solution

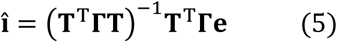

As they are equivalent up to a constant *k*, the solution to constrained linear problem (4a) has the scaled closed-form WLS solution (see Appendix A):

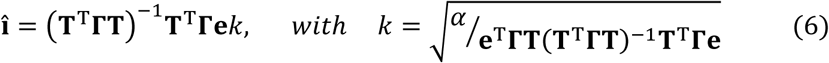

It is easily observed that when non-ROI power bound *α* = **e**^T^**ΓT**(**T**^T^**ΓT**)^-1^**T**^T^**Γe**, *k* = 1 and problems (4a) and (4b) are fully-equivalent.

Problem (4a) above is the constrained maximizing intensity approach (3), but considering that non-ROI electric field energy constraint (3.i) dominates (i.e., *α* is low, and thus total injected current constraint (ii) can be neglected), electric current per electrode bounds (iii) are **ĩ**_**max**_ = *i*_*max*_ and **ĩ**_**min**_ = *–i*_*max*_, i.e. it is allowed that one electrode can inject the total current (constraint (iii) very loose), and **Γ**_**non-roi**_ ≈ **Γ** holds. Thus, we demonstrated that under particular conditions, solution to problem (3) has the closed WLS form of Eq. (6).

If the **Γ**_**non-roi**_ ≈ **Γ** approximation is not considered, solution to problem (4a) becomes *î* = (**T**^T^**Γ**_**non-roi**_**T**)^-1^**T**^T^**Γe**, *k*, with 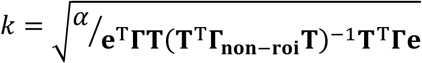. This is not exactly a WLS solution because **Γ**_**non-roi**_ ≠ **Γ**, but extremely similar if ROI ≪ non-ROI, and still a closed-form solution.

If **Γ** is the identity matrix, i.e. there is no integration in Eq. (1) and it describes a pointwise problem, the equivalency between problems (4a) and (4b) still holds, and solution has the “unweighted” LS form: **î** = (**T**^T^**T**)^-1^**T**^T^**e***k*, with 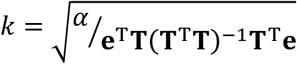.

Overall, if in problem (3), non-ROI energy constraint (3.i) dominates over total injected current bound (3.ii) (meaning that non-ROI energy restriction *α* is low enough such that the solution to problem (3) requires injection of less current than total maximum allowed), the shape of the current injection pattern remains constant (LS or WLS closed formula) regardless the value of α, and α only plays the role of a scaling factor. **One can say the LS or WLS is one “stationary” extreme solution of the constrained directional maximization problem (3) for low α**. This fact is clearly observed in the simulation results (Figs. 1A and 2A).

**Figure 1:**
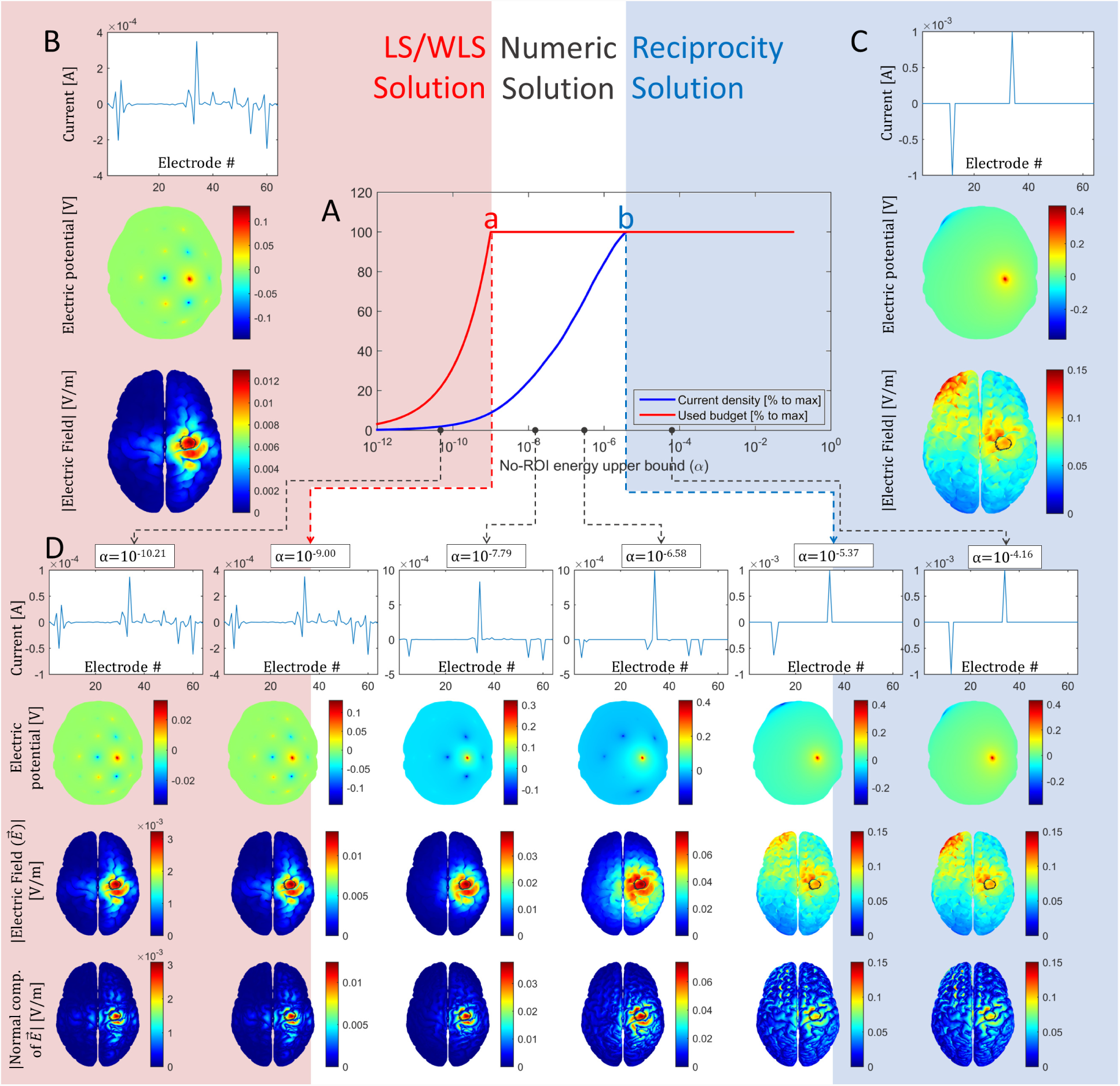
Solutions to the general problem (3) using the trimmed-to-brain electric field transfer matrix. (A) Numerical solutions using the CVX package: percentage of the electric field at the target with respect to the maximum possible (pale blue) and percentage of the total injected current with respect to its maximum budget (red), as a function of the non-ROI power upper bound (a). Each curve has 1000 points equally spaced in logarithmic scale. (B) WLS closed-form solution from Eq. (5) scaled such that the total current injection budget is used. (C) Reciprocity one source - one sink closed-form solution from Eq. (12). (D) Some examples of the numerical solutions. From left to right, starting from low values of *α*, all solutions resemble a WLS closed-form solution (B), except for a scaling constant factor (left side, pale pink zone). When reaching point “a” in plot (A) used budget reaches its upper bound, and the corresponding numerical solution - second column of (D) - is equal to the scaled WLS closed-form solution (B). Between points “a” and “b”, where total allowed budget is exploited, there is a smooth transition between WLS closed-form solution to reciprocity closed-form solution. After point “b” in plot (A), the corresponding numerical solutions depicted in (D) - last two columns - do not change any more and remain equivalent to the closed-form reciprocity one source - one sink solution (C). In subplots (B-D), first row shows the injected current per electrode plot, i.e., the optimal current injection pattern î; second row depicts the electric potential at the scalp as seen from above to spatially visualize the current injection pattern; and third row depicts module of the current density at the brain (i.e., the dose) where the target or ROI is circled in black. In subplot (D), fourth row depicts the absolute value of the normal component of the electric field. Color scale limits are different, increasing from left to right. As this is the trimmed case, similar solutions are obtained if using the current density instead of electric field transfer matrix (see supplementary Fig. S1).

In addition, if total current limit constraint (3.ii) is added to both problems (4.a) and (4.b) the equivalence still holds but for a different value of *α*, which we could not determine analytically. Moreover, if also current limit per electrode (3.iii) is added as well, the equivalence of these problems still holds for a different *α* value. We empirically found these alpha values (see last paragraph of Section 4) and depict corresponding targeting cases in supplementary Figs. S4 and S5. This means that the unified framework of Eq. (3) also covers the “total current constrained” and “fully constrained” LS/WLS solutions (rows 4 and 5 of Table I).

### 3.3 Link between constrained maximizing intensity and reciprocity

The reciprocity theorem coupling TES and EEG states that given a dipole at position 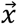 with dipolar moment **d**, the electric potential (Φ) difference between any points *a* and *b* on the scalp can be computed as the dot product:

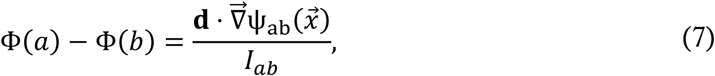

where 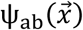 is the resulting potential at location 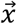 when an electric current *I*_*ab*_ is injected at the arbitrary points *a* and *b* (Malmivuo and Plonsey, 1995; Rush and Driscoll, 1969). In our previous work we demonstrated that if the poles of the EEG forward projection are used for stimulation, the dot product of the electric field and the desired orientation is maximized (Fernández-Corazza et al., 2016). Mathematically,

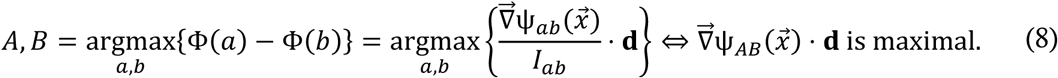

In this work, we go a step further and explicitly link the same reciprocity-based approaches of our previous work with the complete optimization problem (3). For this link to be valid, we assume that *α* is large enough such that total current limit constraint (3.ii) dominates over non-ROI energy limit (3.i) (meaning that non-ROI energy restriction *α* is larger than the maximum possible non-ROI energy, which is achieved in the reciprocity one source-one sink configuration). Note that this assumption results in a similar problem to the simpler maximizing intensity approach of Dmochowski et al 2011 (Eq. (17) in (Dmochowski et al., 2011)) formulated for a pointwise ROI where constraint (3.i) is not considered. Also note that this case is opposite to the extreme case considered in the previous section 3.2, where non-ROI energy limit (3.i) dominates over total current limit constraint (3.ii).

We already mentioned that because e has zeroes in the corresponding non-ROI elements, and assuming the ROI belongs to the brain, the following expression holds: 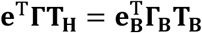, where the sub-index B means the trimmed-to-brain version of the corresponding matrix or vector (recall that **Γ** and **Γ**_**B**_ are the finite element volume diagonal matrices). Setting dipole components to unitless and unit norm magnitude in Eq. (7), by the reciprocity theorem, one can see that the ratio of gradient 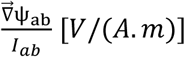 does not depend on injection current being totally defined by geometry an conductivity (as 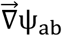 is proportional to *I*_*ab*_). One can more generally obtain from Eq. (7) that elements of TES transfer matrix **T**_**B**_ and EEG Lead Field matrix are related by transposition: **T**_**B**_ **= LF**^T^ (both in [*V*/(*A*. *m*)]), where **LF** is the EEG lead field matrix built as follows: each column of the **LF** matrix corresponds to the electric potential at *L* – 1 electrodes (assuming electrode *L* as the reference) due to a unit dipole at a canonical orientation located at each cortical (or brain) element. Thus, **LF** matrix has size *L* – 1 × 3*N*. The fact that **T**_**B**_ = **L**F^T^ derives from the reciprocity principle in Eq. (7) is well known and demonstrated in the literature (Hallez et al., 2005; Malony et al., 2011; S. Wagner et al., 2016; Weinstein et al., 2000; Wolters et al., 2004), see also more recent discussions in (Dmochowski et al., 2017; Salman et al., 2016). Then,

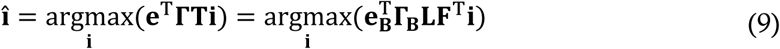

By typical definition of the **LF** matrix, Φ = **LFe**_**B**_, where Φ is the synthetic potential at the electrodes generated by oriented dipoles **e**_**B**_ located at the ROI^3^. Note that the same target vector **e**_**B**_ is considered as EEG sources. We can define a more general potential as Φ = **LF Γ**_***B***_**e**_**B**_, where the effect of **Γ**_**B**_ is just weighting the strength of each dipole that model a volumetric source at the ROI according to the volume of the containing element. Then, expression (9) becomes

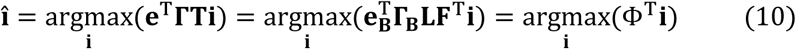

Now, problem (3) is reduced to the constrained linear optimization problem

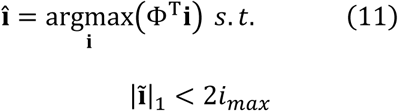

Note that 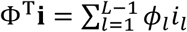, where *ϕ*_*l*_ is the EEG potential at the *l*^th^ electrode. Again, since the functional to maximize is linear, the solution to (11) will lie at the boundary, i.e., when |**ĩ**|_1_ = 2*i*_*max*_. As this problem has a *𝓁*_1_-norm constraint, typical approach until now was using numerical solvers. However, we next show that solution to problem (11) can be found straightforward without the need of numerical solvers:

-Suppose the EEG electric potential Φ generated by artificially placing oriented dipoles at the target has positive and negative values with respect to reference electrode *L*. Then, the sum Φ^**T**^**i** is maximized by the vector **ĩ** that has all zeros except for *i*_*max*_ at the position where Φ is maximum (*ϕ*_*max*_) and *–i*_*max*_ at the position where Φ is minimum (*ϕ*_*min*_), then Φ^T^**î**= *ϕ*_*max*_*i*_*max*_ + (–*i*_*max*_)*ϕ*_*min*_, where *ϕ*_*min*_ is negative. This means that optimal solution is injecting the electric current at the electrodes with maximum and minimum potential (the current at electrode *L* is zero due to Eq. (2)), i.e. injection at the EEG “poles” in a one source - one sink pattern.
-Suppose Φ has all positive values with respect to reference *L*, then Φ^**T**^**i** is maximized by the vector **î** that has all zeros except for *i*_*max*_ at the position where Φ is maximum. Because of Eq. (2), i.e. the total sum of injected current is zero, the sink is the *L* electrode and its strength is *-i*_*max*_.
-Suppose Φ has all negative values, then Φ^T^**i** is maximized by the vector î that has all zeros except for *-i*_*max*_ at the position where Φ is minimum (most negative). Because of Eq. (2), the source is the *L* electrode and its strength is *i*_*max*_.

In all three cases the optimal solution is picking as source and sink the two electrodes with maximum potential difference, i.e., the EEG poles, as described in the one-one approach of Fernandez-Corazza et al 2016 (Fernández-Corazza et al., 2016). Thus, the closed form reciprocity-based one source - one sink solution to problem (11) is:

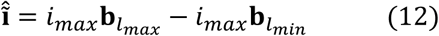

Where **b**_*l*_ is a zero L × 1 vector with a “1” at element “*l*” ^4^, *l*_*max*_ is the electrode with maximum Φ and *l*_*min*_ is the electrode with minimum Φ.

#### 3.3.1 Considering maximum current per electrode limit

If we include maximum current per electrode limit constraint (3.iii), closed-form solutions like (12) can be derived using the same reasoning as described above. For instance, suppose that we set ĩ_max_ *= i*_*max*_/2 and ĩ_min_ *= −i*_*max*_/20. This means that the solution will have at least two sources and twenty sinks. To maximize Φ^T^**i**, the two electrodes with maximum Φ with respect to reference electrode *L* should be selected as sources and the 20 electrodes with minimum Φ with respect to *L* should be selected as sinks. Similarly, it is possible to obtain the “opposite”, “one source-all sinks”, and “10 sources-30 sinks” of Fernandez-Corazza et al 2016 (Fernández-Corazza et al., 2016) by solving problem (3) with corresponding maximum current per electrode constraints imposed by (3.iii)^5^. Thus, these patterns belong to the optimal solution family of the general optimization problem (3) and they are not *ad-hoc* solutions. In the supplementary figure S3, we depict this equivalence with two examples comparing the numerical solutions to (3) considering constraint (3.iii) with the corresponding closed-form reciprocity-based solutions.

## 4 Simulations

In this section we illustrate our analytical findings with simulations on a simple head model based on the ICBM-152 symmetric atlas (Mazziotta et al., 2001).

### 4.1 Simulation framework

We used a head model with four tissues: brain, CSF, skull and scalp based on the ICBM-152 atlas, which is an average of 152 individual heads (Mazziotta et al., 2001). Base-line triangular surfaces were obtained from SPM8 MATLAB package (Friston, 2007) and further refinement, smoothing and tetrahedral meshing was performed using Iso2mesh MATLAB package (Fang and Boas, 2009). The final tetrahedral mesh has ∼1 million elements and ∼150k nodes. We assumed homogeneous and isotropic conductivities for each tissue assigning literature values: 0.3, 0.006, 1.79, and 0.33 S/m for the scalp, skull, CSF, and brain respectively (Baumann et al., 1997; Fernandez-Corazza et al., 2018; Gabriel et al., 1996). The model is completed with 64 pointwise electrodes placed following a subset of standard 10-10 EEG electrode coordinates. Note that we use this simple model to illustrate the theoretical links between different algorithms, and not to evaluate the performance of them. All algorithms can be applied to more complex models with different conductivity values and number of electrodes, as theoretical findings described in previous sections are model-independent.

We selected a part of the M1 cortical region of ∼1.4 cm^3^ as the ROI or target. For each tetrahedral element of the ROI, its centroid was projected to the closest triangular element of the external brain surface and the normal to the cortex vectors of these surface triangles were computed. Desired orientation vector was defined as a weighted by element volume average of these ROI normal vectors. Then, this unique orientation was replicated in each ROI element to form target vector **e**. Note that any other orientation, even arbitrary, can be used instead. Full head transfer matrix **T**_**H**_ was obtained as described in Section 2, and **T**_**B**_ was built by trimming non-brain rows from **T**_**H**_.

### 4.2 Simulations

First, we solved the electric field maximization problem (3) numerically using the CVX package (Grant and Boyd, 2014) for a wide range of *α*, considering total current limit constraint (3.ii) as *i*_*max*_ *=* 1*mA*, and current limit per electrode constraint (3.iii) as *ĩ*_max_ *= i*_*max*_ and *ĩ* _min_ *= −i*_*max*_ (meaning that each electrode can span between *-i*_*max*_ and *i*_*max*_). For each optimal solution of the spanned *α* range, we computed the integral of the electric field over the distributed ROI (i.e., the maximized functional **e**^**T**^**ΓTi** of problem (3)) and the used budget, i.e., the |**ĩ**|_1_. In Figs. 1 and 2, we plot these two values, *each of them normalized to the absolute maximum they can get,* as a function of *α*, for both the trimmed-to-brain (Fig 1) and full-head (Fig 2) transfer matrices ^6^.

**Figure 2:**
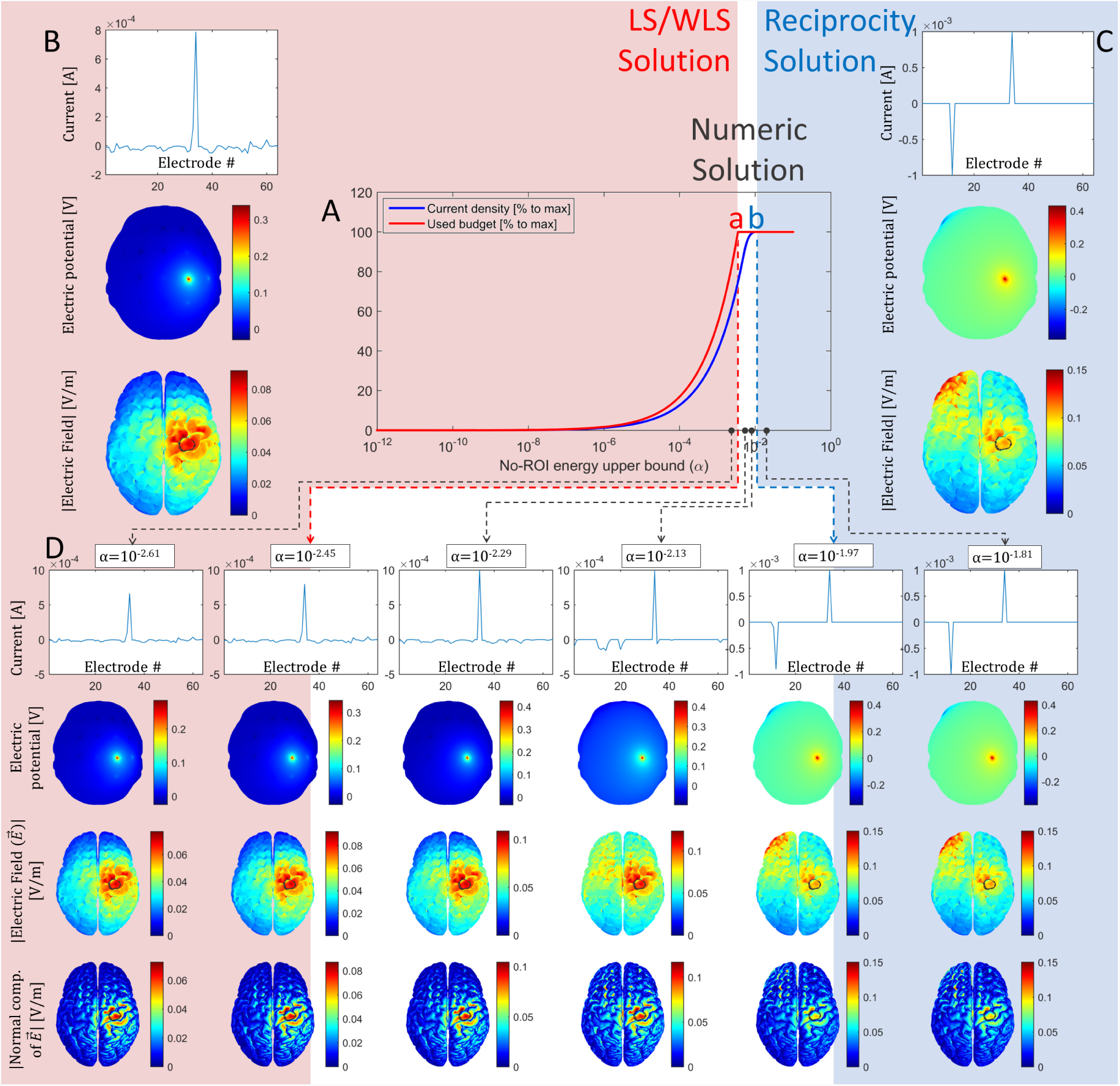
Idem to Fig 1 but using the full-head electric field transfer matrix instead of the trimmed-to-brain transfer matrix. Note that the gap between critical points “a” and “b” is narrower than in Fig. 1, at least in the logarithmic x-axis scaling plot. In this full-head case, different solutions are obtained if using the current density instead of electric field transfer matrix (see supplementary Fig. S2).

In both figures, three different zones can be distinguished, which are separated by colors: the left one (pale pink), where the used budget is less than 100% of the allowed budget, non-ROI energy constraint (3.i) dominates, and total current limit constraint (3.ii) has no influence on the solution; the right zone (pale blue), where the optimal solution remains the same regardless the value of *α*, both the used budget and the maximum current density at ROI saturate, total current limit constraint (3.ii) dominates, and non-ROI energy constraint (3.i) has no influence; and the middle zone (white background), where the current budget saturates but more intense electric field is delivered to the ROI at the expenses of the larger electric field energy allowed at the non-ROI (by increasing *α*). In Figs. 1 and 2, subplots B and C depict the closed form WLS and reciprocity solutions respectively, while subplot D depicts the numerical solution examples for arbitrary selected representative values of *α*.

It is clearly seen that the extremes of the numerical solutions (subplot D) correspond either to a scaled version of the WLS closed-form solution (subplot B) on the left side (as expected due to equivalence between problems (4.a) and (4.b)), or to the reciprocity one source-one sink pattern (subfigure C) on the right side (as expected because Eq. (12) is the solution to problem (9)). In the middle zone, the numerical solution represents a smooth transition from the WLS to reciprocity solution. In the case of using the trimmed-to-brain transfer matrix (Fig. 1), it can be observed that for a particular *α* value, the solution is equivalent to the solution to WLS with 𝓁_1_ constraint (row 4 of Table I) solved using LASSO in Dmochowski et al 2011 (Dmochowski et al., 2011). That the solution is exactly the same is further confirmed in supplementary figure S4. Moreover, many solutions of this middle zone resemble the “ring” patterns of Dmochowski et al 2011 (Dmochowski et al., 2011) and Fernandez-Corazza et al 2016 (Fernández-Corazza et al., 2016). Thus, we empirically found that a series of transitional “ring” solutions (with sequentially increased radius) between LS and the reciprocity one source-one sink patterns are “quasi-optimal” and can be also included as partial solutions of the general optimization problem (3), belonging to the range of the bounding parameter “alpha” in between the WLS and reciprocity-based solutions. A movie showing this transition in great detail can be found as supplementary material.

Comparing critical point “a” in Fig. 1A and Fig. 2A, it is observed that when using the trimmed-to-brain transfer matrix, the WLS solution at point “a” has 10% of the maximum possible intensity at the ROI, the rest of injected currents is shortcut in non-brain tissues, whereas when using the full-head transfer matrix, this intensity (the ROI dose) increases up to 70% of the maximum possible current intensity at the ROI. Also, when using the full-brain transfer matrix (Fig. 2), the middle zone is narrower (at least in the plotted logarithmic scale). Comparing the shapes of the WLS solutions in Figs. 1B and 2B one can see that using the full-head matrix produces a more concentrated solution, as we explicitly include into the optimization problem a demand to avoid “wasting” electric field energy in the rest of the head.

Solving the optimization problem for the current density instead of the electric field, when using the trimmed-to-brain transfer matrix, gives the same results except for a multiplying constant (compare Fig. 1 with supplementary Fig. S1). This is because we are assuming the conductivity values in the domain of the integrals of Eq. (1) are the same: the isotropic and homogeneous conductivity assigned to the brain. When using the full-head transfer matrix, solving the optimization problem for the current density instead of the electric field gives different results because the conductivities differ from tissue to tissue (see supplementary Fig. S2 and related discussion in section 5.5).

As examples, we obtained the numerical solutions of the complete problem (3) in the right pale blue zone (the large *α* values for non-ROI energy bound (3.i)), but setting different current limits per electrode (constraint (3.iii)): as ĩ_max_ *= i*_*max*_ and ĩ _min_ *= −i*_*max*_*/*(*L* - 1), and as ĩ_max_ *= i*_*max*_*/2* and *ĩ*_min_ *= −i*_max_/20. In supplementary Fig. S3 we show that numerical solutions for these two cases are equivalent to the “1 source - (L - 1) sinks” and “2 sources - 20 sinks” closed-form reciprocity-based solutions. Also note that in Figs. 1C, 1D, 2C and 2D, the numerical solutions are equivalent to closed-form one source - one sink reciprocity solution (when ĩ_max_ *= i*_*max*_ and ĩ_min_ *= −i*_*max*_). Thus, we verify that the reciprocity-based patterns involving more than one source or sink, suggested intuitively in (Fernández-Corazza et al., 2016), are optimal solutions to the general problem (3) and not *ad-hoc*.

We also compared the numerical solutions to linear problem (4a) and WLS problem (4b), but explicitly adding the total current limit constraint (3.ii), i.e.:

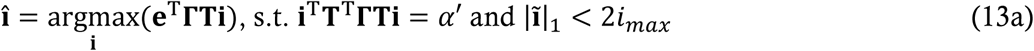

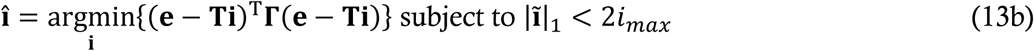

Problem (13b) is a WLS problem with an 𝓁_1_-norm constraint as in Dmochowski et al 2011 (Dmochowski et al., 2011) solved by them using the LASSO solver (Tibshirani, 2011), but here we used CVX for both problems (13a) and (13b). We found that solutions to problems (13a) and (13b) are equivalent for a particular *α′* that we empirically found. Note that in the equivalence of problems 4a and 4b, a formula for *α* was found analytically. This comparison is depicted in figure S4. Moreover, we have further complicated problems (13a) and (13b) by adding a constraint for current per electrode resulting in the following two problems, and also compared the numerical solutions of them:

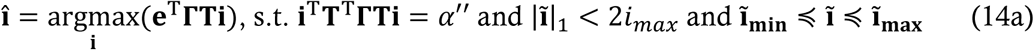

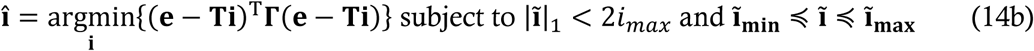

for an example setting ĩ_max_ *= i*_*max*_/2 and *ĩ*_min_ *= −i*_*max*_/20. We again found that both solutions are equivalent for a particular *α″* found empirically. This comparison is shown in supplementary figure S5.

### 4.3 Data and code availability statement

No data was used for the research described in the article. The code supporting the findings of this study are available from the corresponding author upon request.

## 5 Discussion

### 5.1 Links between existing optimization algorithms

We theoretically demonstrated that apparently unrelated LS, WLS, and reciprocity-based solutions all belong to the same family of the general constrained maximizing intensity problem solutions of Eq. (3), constituting a unified approach. This fact was not fully noticed before, and it is the major emphasis of this work. We also found that constrained LS and WLS are also covered by the unified approach and even expanding “ring” configurations are also part of the same family, although these findings were empirical. The transition between WLS or LS to reciprocity-based solutions occurs when increasing (relaxing) the non-ROI energy limit (parameter *α*). LS solutions are recovered at one extreme case (low *α*) and they are more focal and less intense, whereas reciprocity-based solutions belong to the opposite extreme case (large *α*) being less focal and more intense. Ring-shaped patterns occur naturally in the transition zone. Overall, a sliding bar for selecting non-ROI energy upper bound *α* can be included in medical equipment to span the whole range of optimal solutions.

An important demonstration of this work is that the LS or WLS closed-form solution, scaled such that the total current budget is exploited (“scaled LS” in Table I), is also an optimal solution to the general optimization problem (3). This solution equals the numerical solution at critical inflection point marked as “a” in Figs. 1.A, 2.A, S1.A, and S2.A. For lower non-ROI current bounds (*α*), all numerical solutions are identically shaped as the LS or WLS solution but without exploiting the full current injection budget. For larger *α*, the total current limit bound starts shaping the solution producing a smooth transition towards reciprocity solutions.

According to the patterns seen in Figs 3 and 4 of Guler et al 2016 (Guler et al., 2016), numerical solutions resemble the reciprocity “opposite” solutions involving 6-7 electrodes as sources and same number as sinks. Effectively, they adopted a non-ROI energy bound of 10^-6^, which lies in the larger range according to their Fig. 6. The top part of this figure (large *α*) is equivalent to our right “pale blue” area of Figs 1.A, 2.A, S1.A, S2.A, where numerical solutions are equal to reciprocity-based solutions, and bottom part (low *α*) is equivalent to our “left” area where numerical solutions are proportional to closed-form WLS solutions. In Fig. 7 of (Guler et al., 2016), a different (lower) non-ROI energy bound value is used to get similar solutions to the constrained LS solutions by Dmochowski et al 2011 (Dmochowski et al., 2011) and Ruffini et al 2014 (Ruffini et al., 2014).

Quantitative analysis of these algorithms in terms of focality, intensity and other performance metrics for clinical applications is out of the scope of this work, as this was done at length elsewhere (Cancelli et al., 2016; Datta et al., 2013, 2009, Dmochowski et al., 2013, 2011, Fernández-Corazza et al., 2016, 2015; Ruffini et al., 2014; Sven Wagner et al., 2016). The aim of this work is to show the links between many existing methods proposed independently.

Although the analysis done in this work is based in the TES context, the same results can be applied to other techniques. For instance, multi-electrode intracranial electrical stimulation or deep brain stimulation can also benefit from this work (Fernández-Corazza et al., 2017; Guler et al., 2017). The reciprocity principle also holds for the magnetic field and a similar to this TES/EEG duality can be found for TMS/MEG, although a dual TMS/MEG equipment is technically much more complex to build.

### 5.2 WLS vs LS

In Section 3.2, we showed that the integrals in problem (1) derive in a version of WLS where the weights are the volume of each element. However, in most TES optimization approaches using FEM meshes this weighting by element volume was not considered (e.g. (Cancelli et al., 2016; Dmochowski et al., 2011; Fernández-Corazza et al., 2016; Ruffini et al., 2014)) - including our previous work - deriving in unweighted LS. In FEM, the element volumes might vary significantly from each other, thus we believe that the WLS version with the weighting matrix containing the element volumes is more appropriate than the unweighted LS. Of course, additional weighting matrices can also be considered in addition to the volume weighting matrix Γ.

### 5.3 Reciprocity-based algorithms

We found that for a large non-ROI energy bound *α* (loose constraint), and when not considering a current limit per electrode (restriction (3.iii)), solution to problem (3) is equivalent to the reciprocity “one source-one sink” solution. In this solution, the source and sink electrodes are selected directly as the nearest electrodes to the positive and negative forward projection EEG “poles” respectively. Moreover, we found that by setting different current limits per electrode in restriction (3.iii), numerical solutions to problem (3) agreed exactly to other rather intuitive variations of the “opposite” reciprocity solutions described in our previous work (Fernández-Corazza et al., 2016), as shown in supplementary Fig. S3. In such cases, multiple current sources and sinks are selected as close as possible to the EEG forward projection “poles”, with the aim of spreading out the typically undesired electric field concentration due to the sinks.^7^

### 5.4 Full-head or trimmed-to-brain optimization

Typically, trimmed-to-brain (or cortex) transfer matrix was used in the literature (Dmochowski et al., 2011; Guler et al., 2016; Hallez et al., 2005; Malony et al., 2011). In this context, the reciprocal equivalence 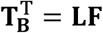 was exploited to compute both LF matrix for EEG source localization and the electric field transfer matrix for TES. This equivalence is obviously also valid for the full-head domain (regardless the conductivity of each tissue), when artificial dipoles are seeded everywhere and not only in the brain to form full LF matrix. We showed that a use of the full-head transfer matrix in TES optimization is beneficial in terms of reducing the ratio of the electric field at the target to the power in the rest of the whole head volume (Fig. 2). When using this approach, the resulting WLS solution is sparser and very similar to the one source - all sinks reciprocity-based solution (see Fig. 2B and Figs 6 and 7 of (Fernández-Corazza et al., 2016)). The use of the trimmed-to-brain transfer matrix can be interpreted as assigning relaxed constraints at the tissues other than the brain. This fact leads to less sparse solutions allowing multiple pairs of nearby sources and sinks shunting each other and located for example on the face and the neck areas far away from the brain, as it is seen in supplementary Fig. S6 when using more electrodes in a more detailed head model (as the one used in our previous work (Fernández-Corazza et al., 2016)). Most of these remote source-sink pairs are likely not contributing into stimulation of the target (see supplementary Fig. S6 and Fig 7 in Guler et al 2016) and, in practice, might lead to the waist of the injection current budget.

### 5.5 Electric field vs current density

The general problem (3) can be solved either using electric field or current in transfer matrices density for dose optimization. What exact physical mechanism causes the wanted and unwanted physiological effects in TES is still in debate to our knowledge.

In most cases there are no significant differences between optimizing electric field or current density, as these physical variables typically (considering isotropy) differ by a scaling constant conductivity factor of approximately a fraction of Siemens per meter (NB: the brain conductivity is in the order of 0.2-0.3 S/m). However, we empirically found that the resulting WLS patterns when using full-head current density transfer matrix are counter-intuitive (see supplementary Fig. S2 left “pale pink” zone examples). This is due to the very low skull conductivity with respect to the other conductivity values of the head. This low value is equivalent to trimming the skull rows from the whole-head transfer matrix (in a simulation experiment not showed here, we empirically forced this situation and results were equivalent to Fig S2 WLS solutions) and thus, it implies a relaxed constraint at the middle skull region. Thus, our recommendation is to optimize electric field when using full head transfer matrix.

### 5.6 Open debates in TES

As we already pointed out, quantitative analysis of each algorithm performance is out of the scope of this work. Each algorithm is optimal in a different sense, for instance, the LS optimization belongs to one end of the optimal solution family, being more focal and less intense and reciprocity-based solutions lie at the other end, being more intense and less focal. We believe that the question of whether focality, total or directional intensity at the ROI should be prioritized for each application is still open. However, considering that the major critique of TES is the low intensity reaching the brain to produce meaningful effects (Horvath et al., 2015), we believe that maximizing the intensity at the target is priority at current stage of TES development. Maximum possible intensity is achieved with reciprocity-based solutions. Note that electric field at point “a” (WLS solution) in Fig. 1 is 10% of the possible maximum intensity at the ROI. To reduce side effects when optimizing for maximum intensity, i.e. undesired intensity at the non-ROI, a possible strategy is to spread the sinks in more electrodes than the sources by setting a lower |**ĩ**_**min**_| than |**ĩ**_**max**_| limit on (3), as in the examples of supplementary Fig. S3 and S5. Once the effects of TES are clearly observed and repeatable, we believe that increasing focality should be the next step in TES optimization evolution.

A question of what orientation is better to target in TES, i.e. which one is more physiologically influential, is still in debate and would depend on the specific application. If pyramidal neurons are the target, perpendicular to cortex stimulation should be preferred, whereas if interneurons synapses are aimed, tangential to cortex stimulation should be more appropriate. The desired orientation can be arbitrarily defined, and all algorithms covered by our unified approach will optimize according to that predefined orientation. In this work, we used normal to cortex orientation to illustrate our results because it is the most commonly used orientation.

Note that while most algorithms optimize directional intensity, in all previous works, the module of the electric field or the current density is depicted. We find this somehow contradictory. That’s why, besides depicting the module, we also depict the normal to cortex component in Figs. 1, 2, S1, S2, S3, S4, and S5. Note that directional optimization methods studied in this work result in much better focality when only considering normal to cortex component instead of module, as some undesired non-ROI “hot-spots” seen when depicting the module are being targeted tangentially to the orientation assumed as physiologically influential.

## 6 Appendix A

Proof that the problem

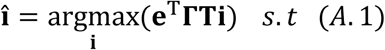

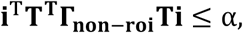

has the solution:

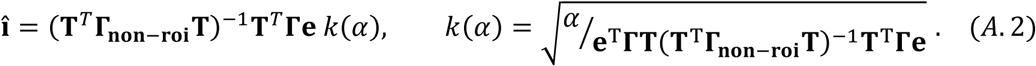

The Lagrangian of the problem is:

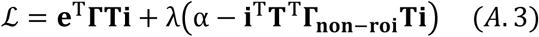

Taking the vector derivative of *ℒ* by *i* and equaling to zero:

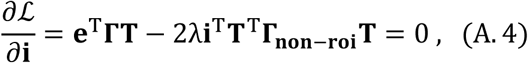

we get:

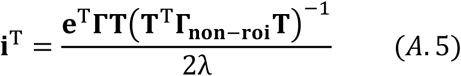

Then, replacing the expression (A.5) into **i**^**T**^**T**^**T**^**Γ**_**non-roi**_**Ti** *=* α, we get (**nr = non - roi**):

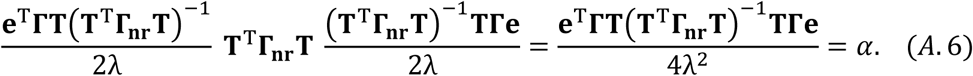

Lastly, one can express:

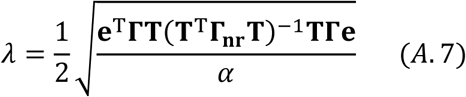

Replacing *λ* in (A.5) with (A.7), one can get the solution (A.2).

Same procedure holds if **Γ**_**non-roi**_ *=* **Γ** as in problems (4.a) and (4.b), resulting in the WLS closed-form solution, which proves equivalence between problems (4.a) and (4.b). As we mention in last paragraphs of Section 4 of this work, this equivalence still holds if we incorporate maximum current limit restriction (3ii) and even more, if we also incorporate current limit per electrode restriction (3.ii) (see examples of Figs. S4 and S5). The difference is that the *α′* and *α″* exact values that make the two problems identical are more difficult to find.

## 7 Acknowledgements

This work was supported by the ANPCyT PICT 2014-1232, UNLP I-209, and CONICET. The work of the second author, ST, was supported in part by the National Institute of Mental Health (Grant R44MH106421).

## 8 Conflicts of interest

Authors declare that they have no conflicts of interest.

## Supplementary material

**Figure S1:**
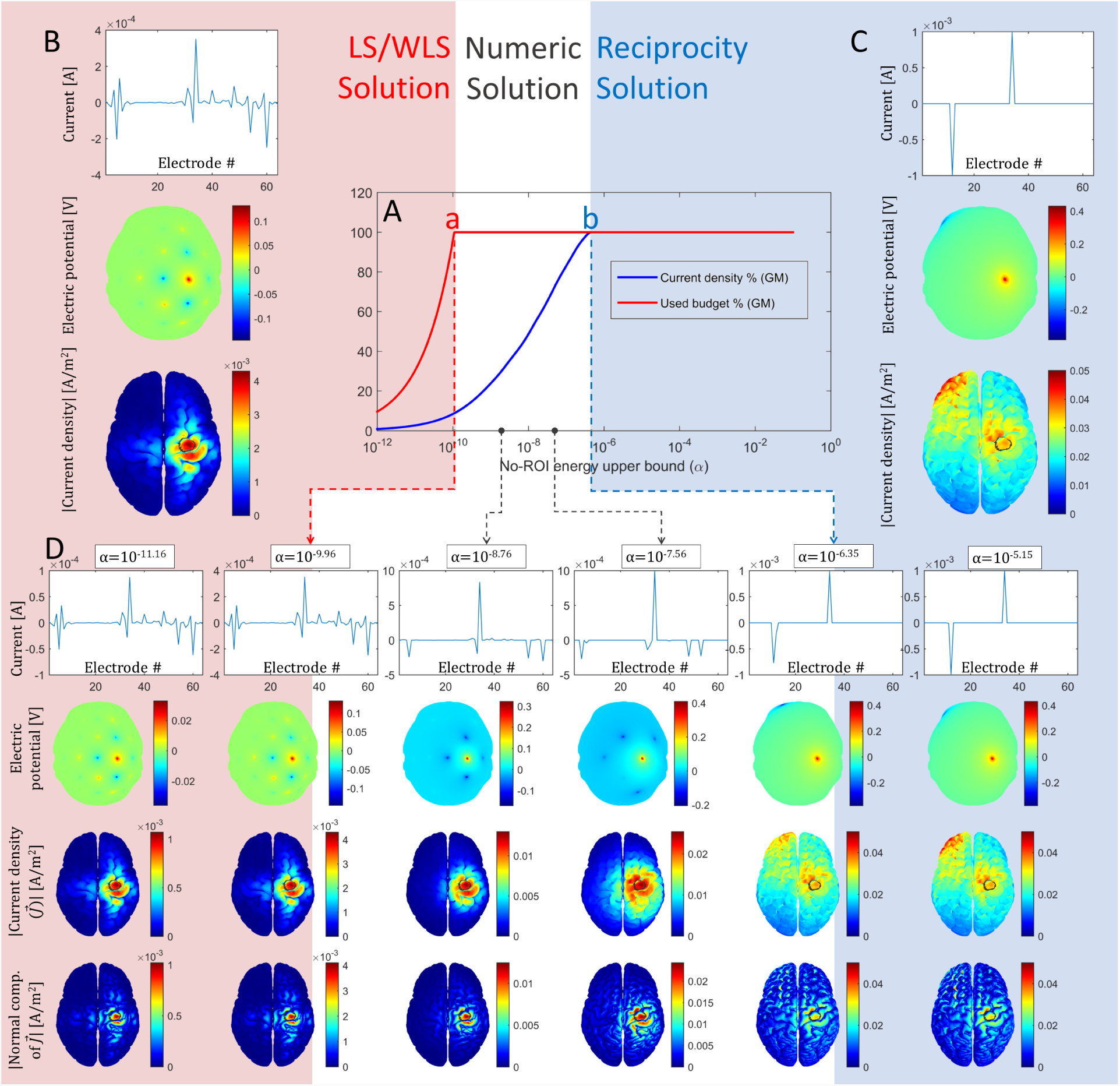
Same as Fig 1 but optimizing the current density instead of the electric field. Note that as here trimmed-to-brain transfer matrix is used, this figure is equivalent to Fig. 1 except for scaling factors.

**Figure S2:**
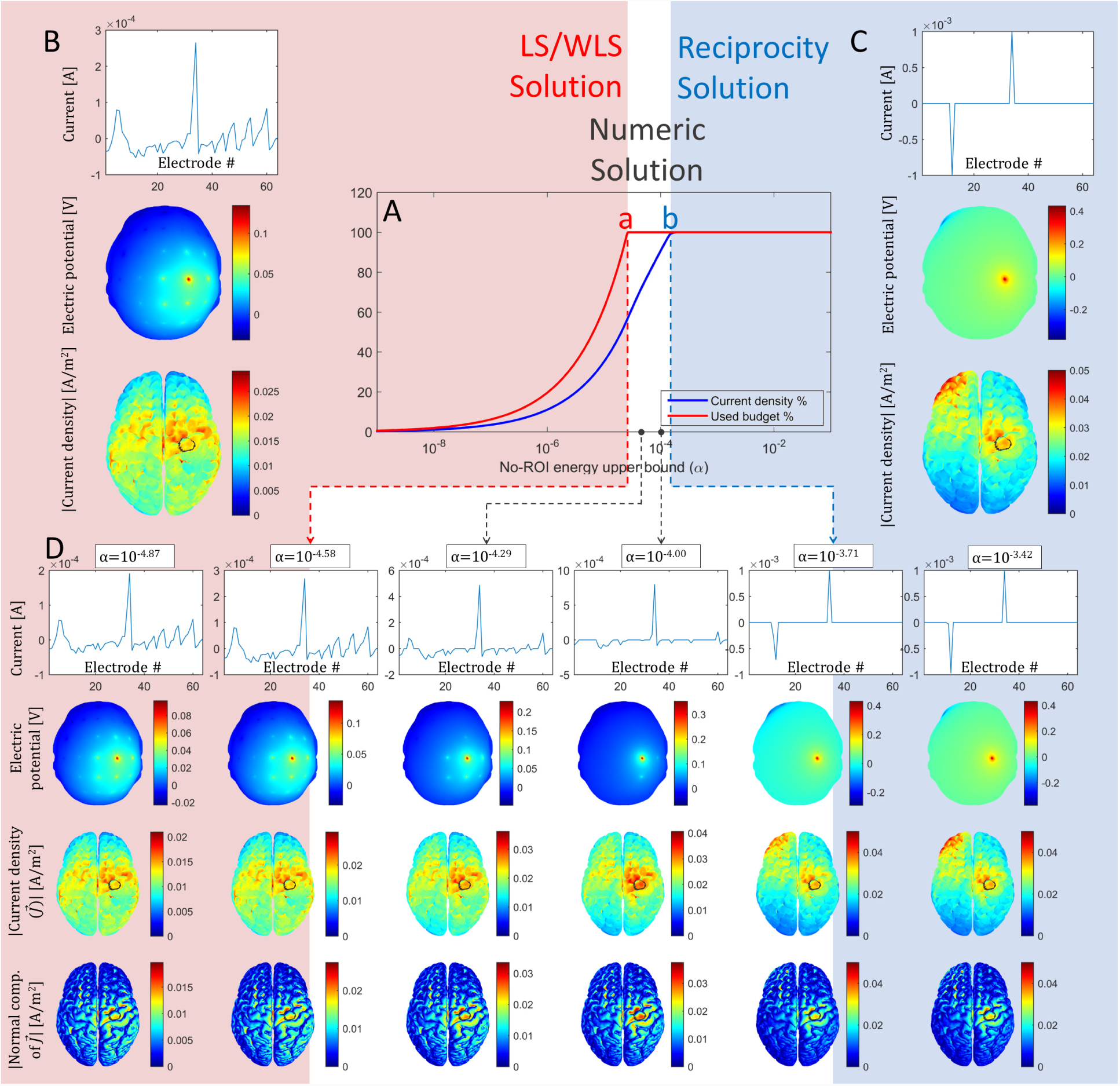
Same as Fig 2 but optimizing the current density instead of the electric field. Note that as here full-head transfer matrix is used, there are different solutions in the WLS pale pink zone compared to Fig 2.

**Figure S3:**
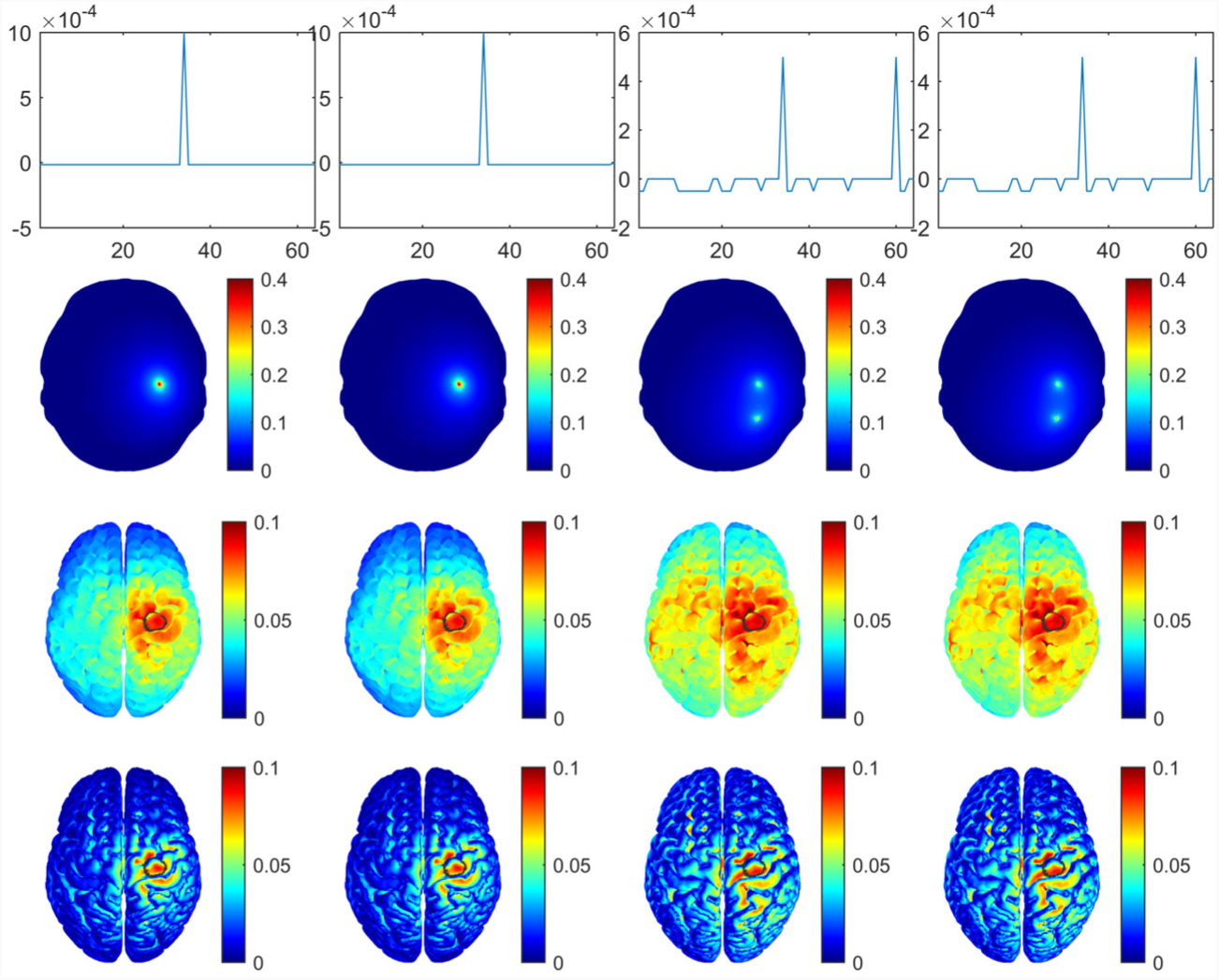
Equivalence between numerical with large α (first and third column) and reciprocity-based closed-form (second and fourth column) solutions when considering limit current per electrode constraint (3.iii) as ĩ_max_ = *i*_max_ and ĩ_min_ = -*i*_max_/(L - 1) (first and second columns); and ĩ_max_ = *i*_max_/2and ĩ_min_ = -*i*_max_/20 (third and fourth columns). First row shows the injected current per electrode [A], i.e., the optimal current injection pattern Î; second row depicts the electric potential [V] at the scalp as seeing form above to spatially visualize the current injection pattern; third row depicts the electric field [V/m] at the brain (i.e., the dose) where the target or ROI is circled in black; and fourth row depicts the absolute value of the normal component of the electric field [V/m].

**Figure S4:**
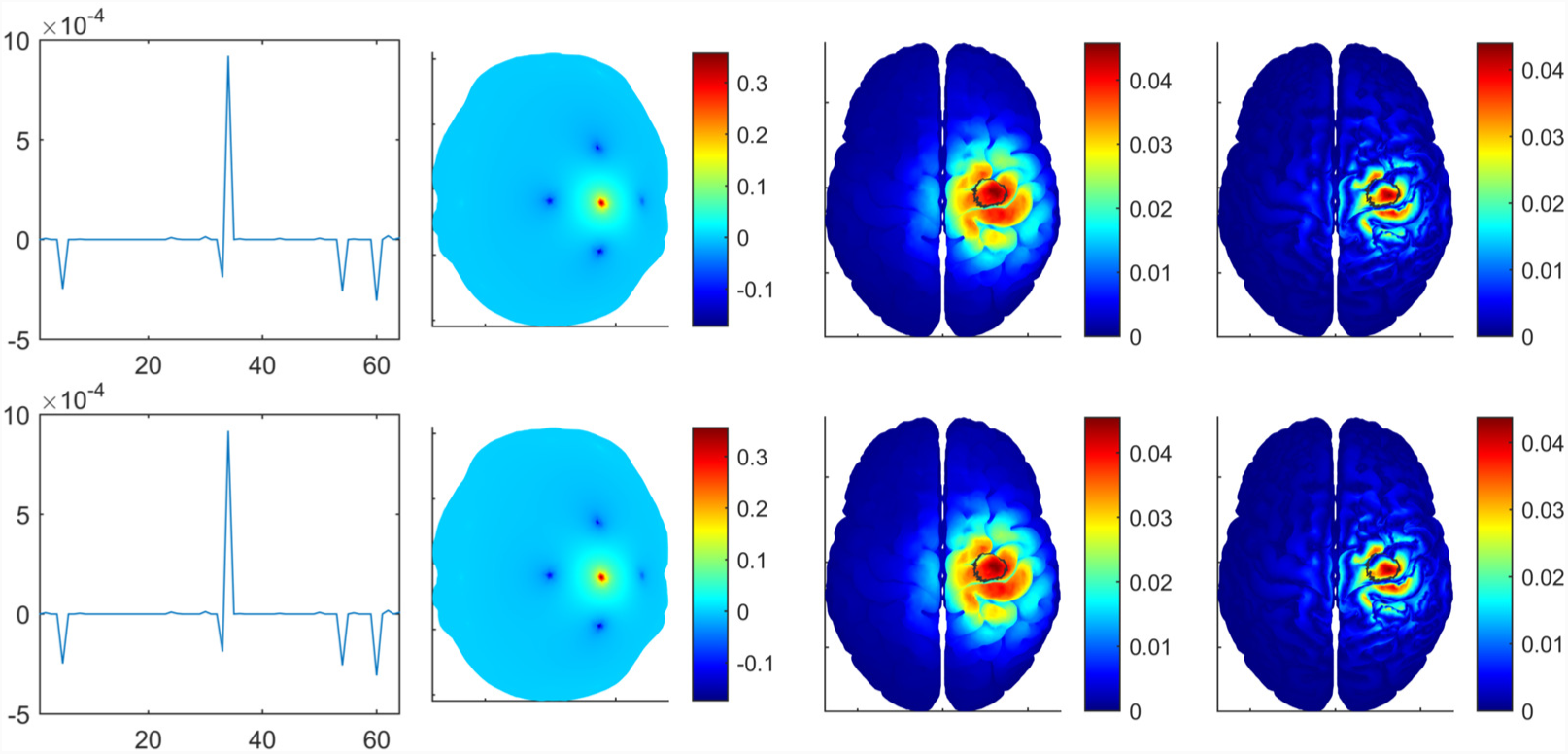
Problems (13.a) and (13.b) are equivalent for a particular α′value. In this example, α′≅ 2.69 × 10^-8^results in the two problems to be equivalent. The first row depicts 𝓁_1_-constrained maximizing intensity solution to problem (13a) using CVX. Second row depicts 𝓁_1_-constrained WLS numerical optimization solution to problem (13b) also using CVX. First column shows the injected current per electrode [A], i.e., the optimal current injection pattern Î; second column depicts the electric potential [V] at the scalp as seeing form above to spatially visualize the current injection pattern; third column depicts the electric field [V/m] at the brain (i.e., the dose) where the target or ROI is circled in black; and fourth column depicts the absolute value of the normal component of the electric field [V/m].

**Figure S5:**
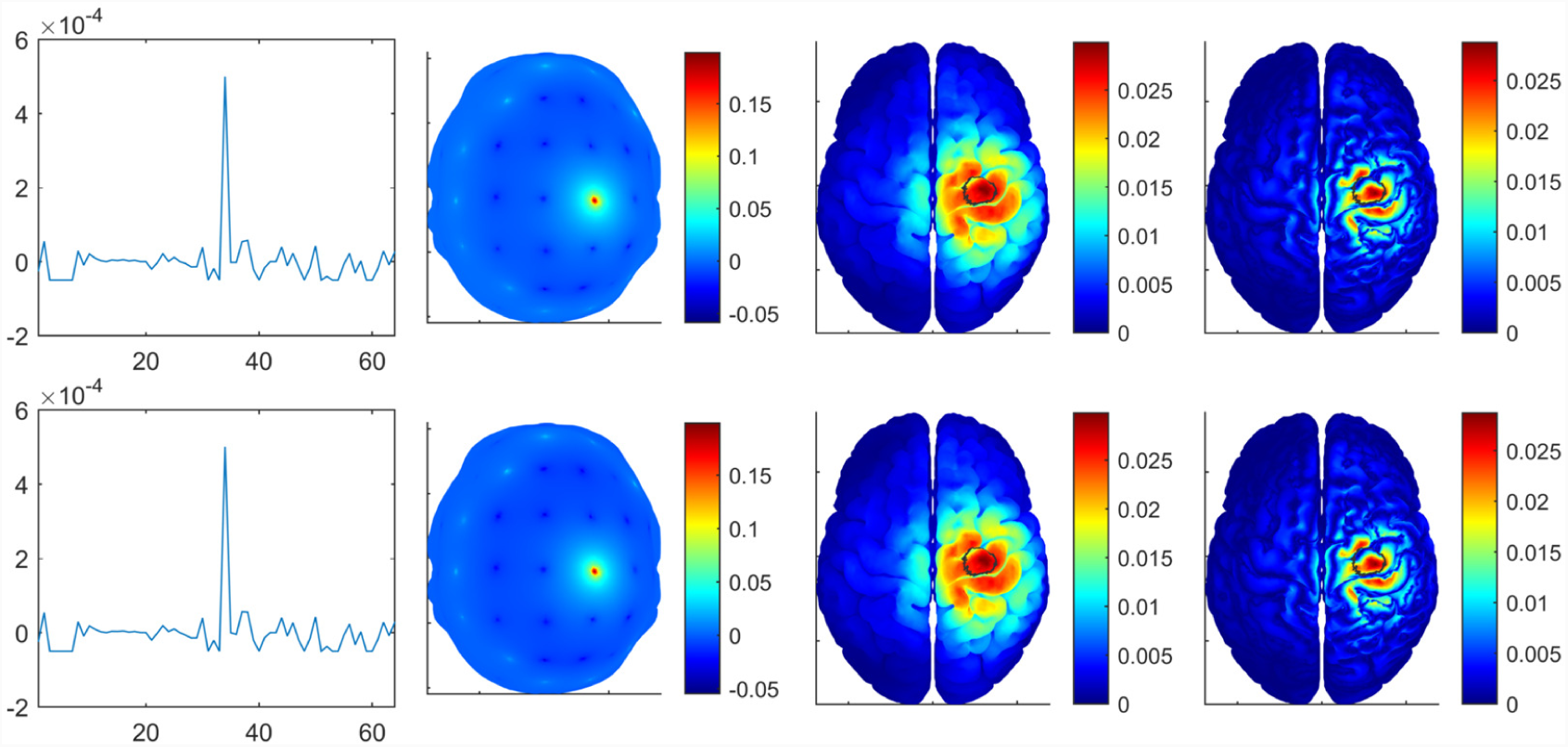
Problems (14.a) and (14.b) are equivalent for a particular α″ value. In this example we set ĩ_max_ = *i*_max_/2 and ĩ_min_ = -*i*_max_/20, i.e. at least two sources and at least 20 sinks, and the resulting α″ value that makes the two problems equivalent was found to be α″ ≅ 1.82 × 10^-8^. The first row depicts total current (𝓁_1_ restriction) and current per electrode constrained maximizing intensity solution to problem (14a) using CVX. Second row depicts total current (𝓁_1_ restriction) and current per electrode constrained WLS numerical optimization solution to problem (14b) also using CVX. First column shows the injected current per electrode [A], i.e., the optimal current injection pattern Î; second column depicts the electric potential [V] at the scalp as seeing form above to spatially visualize the current injection pattern; third column depicts the electric field [V/m] at the brain (i.e., the dose) where the target or ROI is circled in black; and fourth column depicts the absolute value of the normal component of the electric field [V/m].

**Figure S6:**
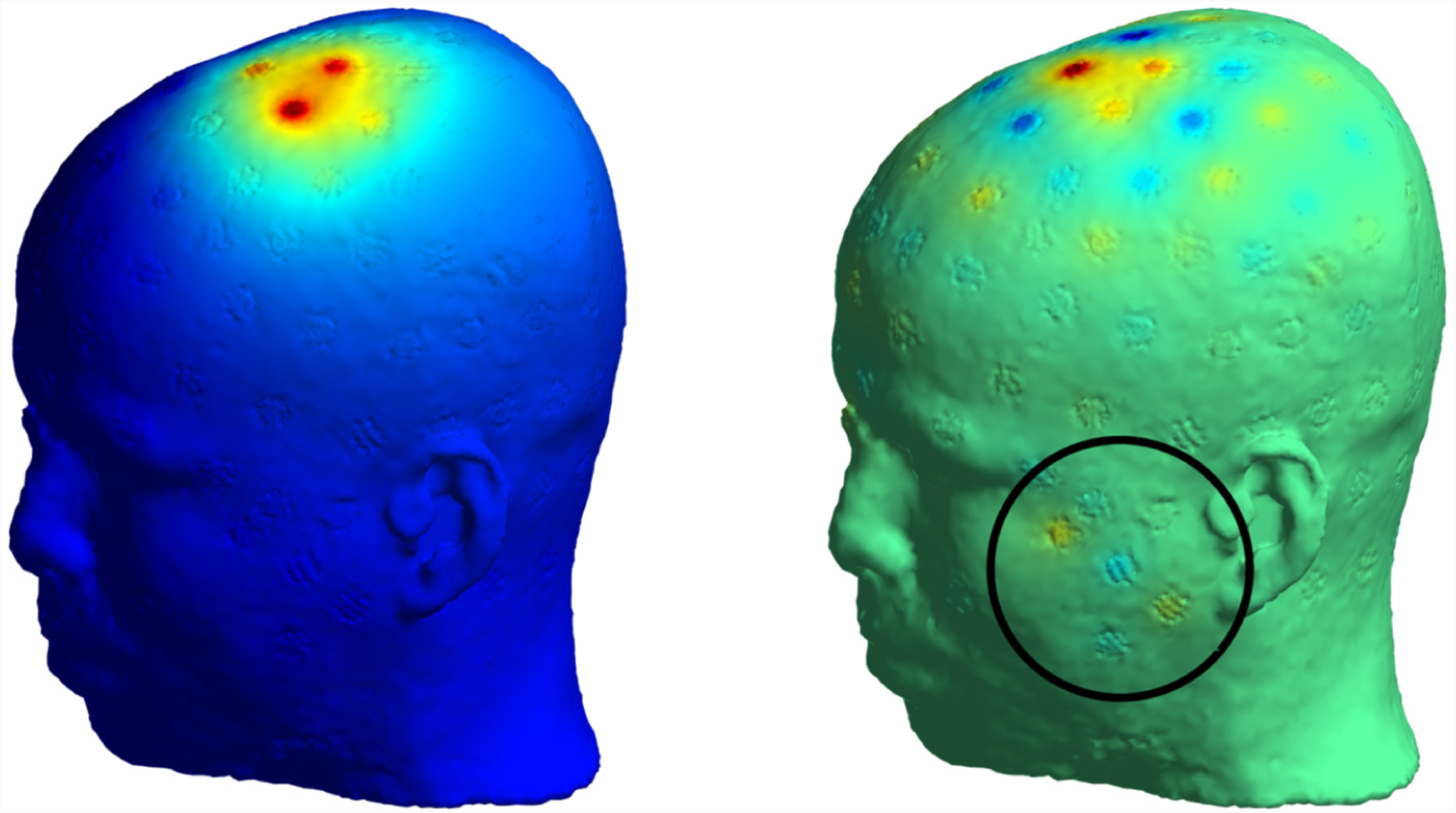
Example of redundant current injection pairs. Same LS closed-form solution for the full-head (left) and trimmed-to-gray matter (right) transfer matrices in a detailed head model (Fernández-Corazza et al., 2016). Example “spurious” current injection pairs appearing in the “trimmed” LS solution are circled in black. As in other figures, the depicted intensity is electric field [V/m] module on the scalp.

Supplementary material: Movie in file named “TES_Unification.avi”.

## Footnotes

1 We use our own notation (the meaning of each symbol/letter can be found in section 3.1) that differs from the authors’ original notation.

2 To be more precise, in (3), **Γ** should be replaced by **Γ**_**roi**_ with non-zero elements only for the elements in the ROI because integral (1) is over the ROI. But, as the elements of **e** corresponding to the non-ROI are set to zero, it is equivalent to use **Γ** or **Γ**_**roi**_ for typical **e**.

3 In order to be consistent with units, **e**_**B**_ has dipole moment units [*A*. *m*].

4 These vectors correspond to the natural basis and are typically notated as **e**_*l*_, for instance in 3D space **e**_1_ = **e**_x_ = [1,0,0]^T^, but as **e** is already used for the desired electric field, we used a different notation **b**_*l*_.

5 To exactly reproduce solutions from Fernandez-Corazza et al 2016 (Fernández-Corazza et al., 2016), Γ in (10) should be set as the identity matrix as in our previous work we didn’t consider integration in the objective function.

6 Note that we know in advance that reciprocity one source - one sink produces the maximum possible current intensity at the ROI. Thus, we precomputed this value and used it to normalize Figs. 1, 2, S1 and S2.

7 For reproducibility, we detail here a minor difference in the target orientation selection between this work and our previous work (Fernández-Corazza et al., 2016). In our previous work, the target orientation was selected as an average of the normal to cortex orientation of each element of the ROI. A better approach to define this orientation is using a weighted average with volume matrix Γ as we did in this work, instead of a simple average as we used in our previous work. In general, the desired orientation selection can be arbitrarily defined, including normal or tangential to brain surface.

## References

Abascal, J.P.J.J., Arridge, S.R., Atkinson, D., Horesh, R., Fabrizi, L., De Lucia, M., Horesh, L., Bayford, R.H., Holder, D.S., Lucia, M. De, Horesh, L., Bayford, R.H., Holder, D.S., 2008. Use of anisotropic modelling in electrical impedance tomography; Description of method and preliminary assessment of utility in imaging brain function in the adult human head. Neuroimage 43, 258–268. doi:10.1016/j.neuroimage.2008.07.023

Antal, A., Keeser, D., Priori, A., Padberg, F., Nitsche, M.A., 2015. Conceptual and Procedural Shortcomings of the Systematic Review “Evidence That Transcranial Direct Current Stimulation (tDCS) Generates Little-to-no Reliable Neurophysiologic Effect Beyond MEP Amplitude Modulation in Healthy Human Subjects: A Systematic R. Brain Stimul. 8, 846–849. doi:10.1016/j.brs.2015.05.010

Barrett, R., Berry, M., Chan, T.F., Demmel, J., Donato, J., Dongarra, J., Eijkhout, V., Pozo, R., Romine, C., der Vorst, H. Van, 1994. Templates for the Solution of Linear Systems: Building Blocks for Iterative Methods, 2nd Edition. SIAM, Philadelphia, PA.

Batsikadze, G., Moliadze, V., Paulus, W., Kuo, M.-F., Nitsche, M.A., 2013. Partially non-linear stimulation intensity-dependent effects of direct current stimulation on motor cortex excitability in humans. J. Physiol. 591, 1987–2000. doi:10.1113/jphysiol.2012.249730

Baumann, S.B., Wozny, D.R., Kelly, S.K., Meno, F.M., 1997. The electrical conductivity of human cerebrospinal fluid at body temperature. IEEE Trans. Biomed. Eng. 44, 220–223. doi:10.1109/10.554770

Berryhill, M.E., Jones, K.T., 2012. tDCS selectively improves working memory in older adults with more education. Neurosci Lett 521, 148–151. doi:10.1016/j.neulet.2012.05.074

Bikson, M., Brunoni, A.R., Charvet, L.E., Clark, V.P., Cohen, L.G., Deng, Z.-D., Dmochowski, J., Edwards, D.J., Frohlich, F., Kappenman, E.S., Lim, K.O., Loo, C., Mantovani, A., McMullen, D.P., Parra, L.C., Pearson, M., Richardson, J.D., Rumsey, J.M., Sehatpour, P., Sommers, D., Unal, G., Wassermann, E.M., Woods, A.J., Lisanby, S.H., 2018. Rigor and reproducibility in research with transcranial electrical stimulation: An NIMH-sponsored workshop. Brain Stimul. 11, 465–480. doi:10.1016/j.brs.2017.12.008

Boggio, P.S., Ferrucci, R., Rigonatti, S.P., Covre, P., Nitsche, M., Pascual-Leone, A., Fregni, F., 2006. Effects of transcranial direct current stimulation on working memory in patients with Parkinson’s disease. J Neurol Sci 249, 31–38.

Boyd, S., Vandenberghe, L., 2004. Convex Optimization. Cambridge Univeristy Press, Cambridge.

Brunoni, A.R., Fregni, F., Priori, A., Ferrucci, R., Boggio, P.S., 2013. Transcranial direct current stimulation: challenges, opportunities and impact on psychiatry and neurorehabilitation. Front. Psychiatry 4. doi:10.3389/fpsyt.2013.00019

Cancelli, A., Cottone, C., Tecchio, F., Truong, D.Q., Dmochowski, J., Bikson, M., 2016. A simple method for EEG guided transcranial electrical stimulation without models. J. Neural Eng. 13, 1–17. doi:10.1088/1741-2560/13/3/036022

Datta, A., Bansal, V., Diaz, J., Patel, J., Reato, D., Bikson, M., 2009. Gyri-precise head model of transcranial direct current stimulation: Improved spatial focality using a ring electrode versus conventional rectangular pad. Brain Stimul 2, 201–207.e1. doi:10.1016/j.brs.2009.03.005

Datta, A., Zhou, X., Su, Y., Parra, L.C., Bikson, M., 2013. Validation of finite element model of transcranial electrical stimulation using scalp potentials: Implications for clinical dose. J Neural Eng 10, 036018. doi:10.1088/1741-2560/10/3/036018

Dmochowski, J.P., Datta, A., Bikson, M., Su, Y., Parra, L.C., 2011. Optimized multi-electrode stimulation increases focality and intensity at target. J Neural Eng 8, 046011. doi:10.1088/1741-2560/8/4/046011

Dmochowski, J.P., Datta, A., Huang, Y., Richardson, J.D., Bikson, M., Fridriksson, J., Parra, L.C., 2013. Targeted transcranial direct current stimulation for rehabilitation after stroke. Neuroimage 75, 12–19. doi:10.1016/j.neuroimage.2013.02.049

Dmochowski, J.P., Koessler, L., Norcia, A.M., Bikson, M., Parra, L.C., 2017. Optimal use of EEG recordings to target active brain areas with transcranial electrical stimulation. Neuroimage 157, 69–80. doi:10.1016/j.neuroimage.2017.05.059

Dutta, A., Dutta, A., 2013. Using electromagnetic reciprocity and magnetic resonance current density imaging to fit multi-electrode montage for non-invasive brain stimulation. Int. IEEE/EMBS Conf. Neural Eng. NER 447–451. doi:10.1109/NER.2013.6695968

Fang, Q., Boas, D.A., 2009. Tetrahedral Mesh Generation from Volumetric Binary and Gray-scale Images, in: Proceedings of the Sixth IEEE International Conference on Symposium on Biomedical Imaging: From Nano to Macro, ISBI’09. IEEE Press, Piscataway, NJ, USA, pp. 1142–1145.

Fernández-Corazza, M., Beltrachini, L., von-Ellenrieder, N., Muravchik, C.H., von Ellenrieder, N., Muravchik, C.H., von-Ellenrieder, N., Muravchik, C.H., 2013. Analysis of parametric estimation of head tissue conductivities using Electrical Impedance Tomography. Biomed Signal Proces 8, 830–837. doi:10.1016/j.bspc.2013.08.003

Fernández-Corazza, M., Collavini, S., Princich, J.P., Kochen, S., Muravchik, C.H., 2017. Electric current distribution in pre-surgical neurostimulation using stereo-EEG electrodes, in: Libro de Resúmenes XXI Congreso Argentino de Bioingeniería SABI2017. Ciudad de Córdoba, Argentina, p. 56.

Fernández-Corazza, M., Turovets, S., Luu, P., Anderson, E., Tucker, D., 2016. Transcranial Electrical Neuromodulation Based on the Reciprocity Principle. Front. Psychiatry 7. doi:10.3389/fpsyt.2016.00087

Fernandez-Corazza, M., Turovets, S., Luu, P., Muravchik, C., Tucker, D., 2017. Use of reciprocity in TES: ways of choosing the current injection electrodes. Brain Stimul. 10, 401–402. doi:10.1016/j.brs.2017.01.191

Fernandez-Corazza, M., Turovets, S., Luu, P., Price, N., Muravchik, C.H., Tucker, D., 2018. Skull modeling effects in conductivity estimates using parametric electrical impedance tomography. IEEE Trans. Biomed. Eng. 65, 1785–1797. doi:10.1109/TBME.2017.2777143

Fernández-Corazza, M., Turovets, S., Luu, P., Tucker, D., 2015. Optimization in transcranial electrical neuromodulation based on the reciprocity principle. Brain Stimul. 8, 403. doi:http://dx.doi.org/10.1016/j.brs.2015.01.286

Frank, E., 1952. Electric Potential Produced by Two Point Current Sources in a Homogeneous Conducting Sphere. J. Appl. Phys. 23, 1225–1228.

Friston, K., 2007. Statistical Parametric Mapping: The Analysis of Functional Brain Images, 1st Editio. ed. Elsevier/Academic Press, Amsterdam.

Gabriel, S., Lau, R.W., Gabriel, C., 1996. The dielectric properties of biological tissues: II. Measurements in the frequency range 10 Hz to 20 GHz. Phys. Med. Biol. 41, 2251–69.

Goncalves, S.I., de Munck, J.C., Verbunt, J.P. a., Bijma, F., Heethaar, R.M., Lopes da Silva, F., 2003. In vivo measurement of the brain and skull resistivities using an eit-based method and realistic models for the head. IEEE Trans. Biomed. Eng. 50, 754–767. doi:10.1109/TBME.2003.812164

Grant, M., Boyd, S., 2014. CVX: Matlab software for disciplined convex programming, version 2.1. [WWW Document]. URL http://cvxr.com/cvx

Guhathakurta, D., Dutta, A., 2016. Computational pipeline for NIRS-EEG joint imaging of tDCS-evoked cerebral responses-An application in ischemic stroke. Front. Neurosci. 10. doi:10.3389/fnins.2016.00261

Guler, S., Dannhauer, M., Erem, B., Macleod, R., Tucker, D., Turovets, S., Luu, P., Erdogmus, D., Brooks, D.H., 2016. Optimization of focality and direction in dense electrode array transcranial direct current stimulation (tDCS). J. Neural Eng. 13, 1–14. doi:10.1088/1741-2560/13/3/036020

Guler, S., Dannhauer, M., Roig-Solvas, B., Gkogkidis, A., MacLeod, R., Ball, T., Ojemann, J.G., Brooks, D.H., 2017. An algorithm to optimize ECoG stimulus current patterns while constraining local current density across the entire brain. Brain Stimul. 10, e38. doi:10.1016/j.brs.2017.04.065

Hallez, H., Vanrumste, B., Hese, P. Van, D’Asseler, Y., Lemahieu, I., Walle, R. Van de, 2005. A finite difference method with reciprocity used to incorporate anisotropy in electroencephalogram dipole source localization. Phys. Med. Biol. 50, 3787–3806. doi:10.1088/0031-9155/50/16/009

Horvath, J.C., Carter, O., Forte, J.D., 2014. Transcranial direct current stimulation: five important issues we aren’t discussing (but probably should be). Front. Syst. Neurosci. 8. doi:10.3389/fnsys.2014.00002

Horvath, J.C., Forte, J.D., Carter, O., 2015. Evidence that transcranial direct current stimulation (tDCS) generates little-to-no reliable neurophysiologic effect beyond MEP amplitude modulation in healthy human subjects: A systematic review. Neuropsychologia 66, 213–236. doi:10.1016/j.neuropsychologia.2014.11.021

Hyvönen, N., 2004. Complete Electrode Model of Electrical Impedance Tomography: Approximation Properties and Characterization of Inclusions. SIAM J. Appl. Math. 64, 902–931. doi:10.1137/S0036139903423303

Ilmoniemi, R.J., Ruohonen, J., Karhu, J., 1999. Transcranial magnetic stimulation-a new tool for functional imaging of the brain. Crit. Rev. Biomed. Eng. 27, 241–84.

Jackson, J.D., 1975. Classical Electrodynamics Second Edition. John Wiley & Sons, New York.

Kalu, U.G., Sexton, C.E., Loo, C.K., Ebmeier, K.P., 2012. Transcranial direct current stimulation in the treatment of major depression: a meta-analysis. Psychol. Med. 42, 1791–1800. doi:10.1017/S0033291711003059

Kwon, Y.W., Bang, H., 2000. The Finite Element Method Using MATLAB (2Nd Ed.). CRC Press, Inc., Boca Raton, FL, USA.

Laakso, I., Tanaka, S., Mikkonen, M., Koyama, S., Sadato, N., Hirata, A., 2016. Electric fields of motor and frontal tDCS in a standard brain space: A computer simulation study. Neuroimage 137, 140–151. doi:10.1016/j.neuroimage.2016.05.032

Lang, N., Siebner, H.R., Ward, N.S., Lee, L., Nitsche, M.A., Paulus, W., Rothwell, J.C., Lemon, R.N., Frackowiak, R.S., 2005. How does transcranial DC stimulation of the primary motor cortex alter regional neuronal activity in the human brain? Eur J Neurosci 22, 495–504.

Lionheart, W., Polydorides, N., Borsic, a., 2004. The Reconstruction Problem, Electrical Impedance Tomography: Methods, History and Applications. Inst. Phys. doi:10.1118/1.1995712

Malmivuo, J., Plonsey, R., 1995. Bioelectromagnetism: Principles and Applications of Bioelectric and Biomagnetic Fields. Oxford University Press.

Malony, A.D.A.D., Salman, A., Turovets, S., Tucker, D., Volkov, V., Li, K., Song, J.E.J.E., Biersdorff, S., Davey, C., Hoge, C., Hammond, D., 2011. Computational modeling of human head electromagnetics for source localization of milliscale brain dynamics. Stud. Health Technol. Inform. 163, 329–35. doi:10.3233/978-1-60750-706-2-329

Mazziotta, J.C., Toga, A., Evans, A., Fox, P., Lancaster, J.L., Zilles, K., Woods, R., Paus, T., Simpson, G., Pike, B., Holmes, C., Collins, L., Thompson, P., MacDonald, D., Iacoboni, M., Schormann, T., Amunts, K., Palomero-Gallagher, N., Geyer, S., Parsons, L., Narr, K., Kabani, N., Goualher, G. Le, Boomsma, D., Cannon, T., Kawashima, R., Mazoyer, B., 2001. A probabilistic atlas and reference system for the human brain: International Consortium for Brain Mapping (ICBM). Philos. Trans. R. Soc. Lond. B Biol. Sci. 356, 1293–1322.

Mori, F., Codecà, C., Kusayanagi, H., Monteleone, F., Buttari, F., Fiore, S., Bernardi, G., Koch, G., Centonze, D., 2010. Effects of Anodal Transcranial Direct Current Stimulation on Chronic Neuropathic Pain in Patients With Multiple Sclerosis. J. Pain 11, 436–442. doi:10.1016/j.jpain.2009.08.011

Nitsche, M.A., Paulus, W., 2000. Excitability changes induced in the human motor cortex by weak transcranial direct current stimulation. J. Physiol. 527, 633–639. doi:10.1111/j.1469-7793.2000.t01-1-00633.x

Nitsche, M.A., Schauenburg, A., Lang, N., Liebetanz, D., Exner, C., Paulus, W., Tergau, F., 2003. Facilitation of implicit motor learning by weak transcranial direct current stimulation of the primary motor cortex in the human. J. Cogn. Neurosci. 15, 619–626. doi:10.1162/089892903321662994

Otal, B., Dutta, A., Foerster, Á., Ripolles, O., Kuceyeski, A., Miranda, P.C., Edwards, D.J., Ilic, T. V, Nitsche, M.A., Ruffini, G., 2016. Opportunities for Guided Multichannel Non-invasive transcranial current stimulation in Poststroke rehabilitation 7. doi:10.3389/fneur.2016.00021

Priori, A., 2003. Brain polarization in humans: A reappraisal of an old tool for prolonged non-invasive modulation of brain excitability. Clin Neurophysiol 114, 589–595.

Priori, A., Berardelli, A., Rona, S., Accornero, N., Manfredi, M., 1998. Polarization of the human motor cortex through the scalp. Neuroreport 9, 2257–2260. doi:10.1097/00001756-199807130-00020

Ruffini, G., Fox, M.D., Ripolles, O., Miranda, P.C., Pascual-Leone, A., 2014. Optimization of multifocal transcranial current stimulation for weighted cortical pattern targeting from realistic modeling of electric fields. Neuroimage 89, 216–225. doi:10.1016/j.neuroimage.2013.12.002

Rush, S., Driscoll, D.A., 1969. EEG Electrode Sensitivity-An Application of Reciprocity. IEEE Trans. Biomed. Eng. BME-16, 15–22. doi:10.1109/TBME.1969.4502598

Sadleir, R., Vannorsdall, T.D., Schretlen, D.J., Gordon, B., 2012. Target Optimization in tDCS. Front. Psychiatry 3.

Salman, A., Malony, A., Turovets, S., Volkov, V., Ozog, D., Tucker, D., 2016. Concurrency in electrical neuroinformatics: parallel computation for studying the volume conduction of brain electrical fields in human head tissues. Concurr. Comput. Pract. Exp. 28, 2213–2236. doi:10.1002/cpe.3510

Schlaug, G., Renga, V., Nair, D., 2008. Transcranial direct current stimulation in stroke recovery. Arch Neurol 65, 1571–1576. doi:10.1001/archneur.65.12.1571

Silvester, P.P., Ferrari, R.L., 1994. Finite Elements for Electrical Engineers. Cambridge University Press, Cambridge.

Tibshirani, R., 2011. Regression shrinkage and selection via the lasso: a retrospective. J. R. Stat. Soc. Ser. B (Statistical Methodol. 73, 273–282. doi:10.1111/j.1467-9868.2011.00771.x

Turovets, S., Volkov, V., Zherdetsky, A., Prakonina, A., Malony, A.D., 2014. A 3D Finite-Difference BiCG Iterative Solver with the Fourier-Jacobi Preconditioner for the Anisotropic EIT/EEG Forward Problem. Comput. Math. Methods Med. 2014, 1–12. doi:10.1155/2014/426902

van der Vorst, H.A., 1992. Bi-CGSTAB: A Fast and Smoothly Converging Variant of Bi-CG for the Solution of Nonsymmetric Linear Systems. SIAM J. Sci. Stat. Comput. 13, 631–644. doi:10.1137/0913035

Vauhkonen, P.J., Vauhkonen, M., Savolainen, T., Kaipio, J.P., 1999. Three-dimensional electrical impedance tomography based on the complete electrode model. IEEE Trans. Biomed. Eng. 46, 1150–1160. doi:10.1109/10.784147

Wagner, S., Burger, M., Wolters, C.H., 2016. An Optimization Approach for Well-Targeted Transcranial Direct Current Stimulation. SIAM J. Appl. Math. 76, 2154–2174. doi:10.1137/15M1026481

Wagner, S., Lucka, F., Vorwerk, J., Herrmann, C.S., Nolte, G., Burger, M., Wolters, C.H., 2016. Using reciprocity for relating the simulation of transcranial current stimulation to the EEG forward problem. Neuroimage 140, 163–173. doi:10.1016/j.neuroimage.2016.04.005

Wang, P., Li, H., Xie, L., Sun, Y., 2009. The Implementation of FEM and RBF Neural Network in EIT. 2009 Second Int. Conf. Intell. Networks Intell. Syst. 66–69. doi:10.1109/ICINIS.2009.26

Weinstein, D., Zhukov, L., Johnson, C., 2000. Lead-field bases for electroencephalography source imaging. Ann. Biomed. Eng. 28, 1059–1065. doi:10.1114/1.1310220

Wiethoff, S., Hamada, M., Rothwell, J.C., 2014. Variability in Response to Transcranial Direct Current Stimulation of the Motor Cortex. Brain Stimul. 7, 468–475. doi:10.1016/j.brs.2014.02.003

Windhoff, M., Opitz, A., Thielscher, A., 2013. Electric field calculations in brain stimulation based on finite elements: An optimized processing pipeline for the generation and usage of accurate individual head models. Hum. Brain Mapp. 34, 923–935. doi:10.1002/hbm.21479

Wolters, C.H., Grasedyck, L., Hackbusch, W., 2004. Efficient computation of lead field bases and influence matrix for the FEM-based EEG and MEG inverse problem. Inverse Probl 20, 1099. doi:10.1088/0266-5611/20/4/007

Yook, S.-W., Park, S.-H., Seo, J.-H., Kim, S.-J., Ko, M.-H., 2011. Suppression of Seizure by Cathodal Transcranial Direct Current Stimulation in an Epileptic Patient - A Case Report -. Ann Rehabil Med 35, 579–582. doi:10.5535/arm.2011.35.4.579

